# A micropeptide component of the OGDH complex modulates ATP production in the TCA cycle

**DOI:** 10.64898/2026.01.12.699137

**Authors:** Xinqiang Yin, Weiyan Qi, Lifen Lin, Huilin Zheng, Quan Chen, Guoxiang Zhao, Guoqiang Shao, Yifan Wu, Jialiang Hu, Hanmei Xu

**Affiliations:** The Engineering Research Center of Synthetic Polypeptide Drug Discovery and Evaluation, Jiangsu Province, China Pharmaceutical University, Nanjing 210009, P.R. China; School of Basic Medicine and Forensic Medicine, North Sichuan Medical College, Nanchong, 673000, P.R. China; Department of Nuclear Medicine, Nanjing First Hospital, Nanjing Medical University, Nanjing 210009, P.R. China; Mudi Meng Honors College, China Pharmaceutical University, Nanjing 211198, P.R. China; State Key Laboratory of Natural Medicines, Ministry of Education, China Pharmaceutical University, Nanjing 210009, P.R. China

**Author notes:** Joint International Research Laboratory of Target Discovery and New Drug Innovation, MOE. These authors contributed equally to this work (Xinqiang Yin, Weiyan Qi).

**Keywords:** Micropeptde APPM, Mitochondria, Tricarboxylic acid cycle, OGDH, Heart failure, Bioenergetic disorders

## Abstract

Since its elucidation, the tricarboxylic acid (TCA) cycle has been regarded as a fully defined enzymatic pathway. Here, we report the discovery of a previously unrecognized micropeptide component of this central metabolic system. Through analyzing ribosome profiling data from the mouse sORFs.org database, we discovered APPM (ATP production–promoting micropeptide), a 29–amino acid peptide encoded by a non-canonical small open reading frame within the mouse *Ctsb* gene. APPM localizes to the mitochondrial matrix and inner membrane, where it directly binds to the E1 subunit (OGDH) of the α-ketoglutarate dehydrogenase complex (OGDHc). This interaction modulates OGDHc enzymatic activity, thereby regulating NADH and ATP production. We identified tyrosine-13 as the critical residue mediating the APPM-OGDH interaction and functional activation. Exogenous APPM administration increased ATP levels *in vivo* and, in a mouse model of heart failure, restored cardiac function, outperforming the reference drug trimetazidine. Collectively, our findings identify APPM as the first micropeptide regulator of the OGDH complex, expanding the known repertoire of the TCA cycle and presenting a potential therapeutic strategy for bioenergetic disorders.

---

Mitochondria are the central powerhouses of eukaryotic cells, generating ATP through oxidative phosphorylation to sustain cellular metabolism and physiological functions. At the core of this process lies the tricarboxylic acid (TCA) cycle, a central metabolic pathway responsible for the oxidation of carbohydrates, lipids, and amino acids, while also supplying crucial biosynthetic precursors. Dysregulation of the TCA cycle is a hallmark of mitochondrial dysfunction and underlies a broad spectrum of human diseases, including neurodegenerative, cardiovascular, and metabolic disorders^1–3^.

Within the TCA cycle, the α-ketoglutarate dehydrogenase complex (OGDHc) functions as a critical rate-limiting enzyme catalyzing the conversion of α-ketoglutarate (α-KG) to succinyl-CoA, while reducing NAD⁺ to NADH, a key electron donor for the electron transport chain^4^. The activity of OGDHc is finely tuned by metabolic feedback inhibition via its products, succinyl-CoA and NADH, and allosteric activation by ADP and calcium ions^5^. Consequently, impaired OGDHc function compromises ATP synthesis, alters mitochondrial dynamics, and has been linked to severe metabolic abnormalities, including developmental delays, movement disorders, and lactic acidosis^6–10^.

Since the TCA cycle was elucidated by Hans Adolf Krebs, —earning the 1953 Nobel Prize in Physiology or Medicine, — its core enzymatic machinery has been considered fully mapped and remained remarkably conserved. Subsequent research has primarily extended this framework, focusing on regulatory mechanisms such as post-translational modifications of OGDHc components (e.g., phosphorylation^11^, SUMOylation^12^), interactions with chaperones like HSP60^13^, and modulation by non-coding RNAs^14^. Despite these advances, no additional core protein constituents of the TCA cycle have been identified for more than seven decades.

In parallel, the recent emergence of micropeptides—small proteins typically fewer than 100 amino acids encoded by non-canonical small open reading frames (sORFs)—has unveiled a hidden layer of regulatory biology. These micropeptides have been shown to play critical roles in diverse cellular processes, including ion channel function, cellular signaling, and RNA metabolism^15–19^. Our previous work contributed to this field by identifying micropeptides such as MIAC, which suppresses tumor growth in head and neck squamous cell carcinoma and renal cell carcinoma by interacting with AQP2^20,21^. Such findings, together with others, have led to the reclassification of certain non-coding RNAs as “potential-coding RNAs” (pcRNAs)^22^.

Notably, over 20 mitochondrial-targeted micropeptides have been identified, regulating processes ranging from respiratory complex assembly and mitochondrial biogenesis to apoptosis^23–30^. While some mitochondrial micropeptides indirectly influence ATP levels^31–37^, none have been reported to directly bind and modulate the core enzymes of the TCA cycle to regulate ATP generation.

In this study, we systematically screened an alternative open reading frame (alt-ORF) within the mouse *Ctsb* gene using ribosome profiling data from the sORFs.org database^38,39^. Through a combination of *in vivo* and *in vitro* experiments, multi-omics analysis, and molecular biology techniques, we validated and characterized a micropeptide named APPM (ATP production–promoting micropeptide) which was encoded by this alt-ORF. We further elucidated its molecular mechanism of action and biological significance, uncovering a direct regulatory link between a micropeptide and the core machinery of the TCA cycle.

## Results

### Screening of the sORF encoding APPM

To systematically identify functional micropeptides, a stringent bioinformatic screening of mouse ribosome profiling data was performed and followed by experimental validation. Ribosome profiling data were retrieved from the sORFs.org database, comprising 267,362 entries derived from 16 independent mouse studies. After filtering out redundant data, 71,783 sORF entries remained.

From these, top 10 sORFs derived from the sense strand, without splicing, and with the highest PhyloP scores were selected. Based on the gene localization, upstream/downstream sequence information, ribosome binding sequences, as well as mutation sites and phenotypic data from dbSNP within these sORFs, 5 sORFs were ultimately selected for further study (Fig. 1A). *sORF-Fc* fusion genes (with the Fc-encoding sequence inserted before the stop codon of the sORF) were constructed. The fusion genes were cloned into the pcDNA3.0 plasmid vector and transfected into CHO cells. Using an anti-Fc antibody, the APPM-Fc fusion product was detected in total protein extracts from CHO cells (Fig. 1B). This sORF is a 90-nucleotide alternative open reading frame (alt-ORF) located within the exonic and intronic regions of the CTSB protein-coding gene (Fig. 1C, D).

**Figure 1.**
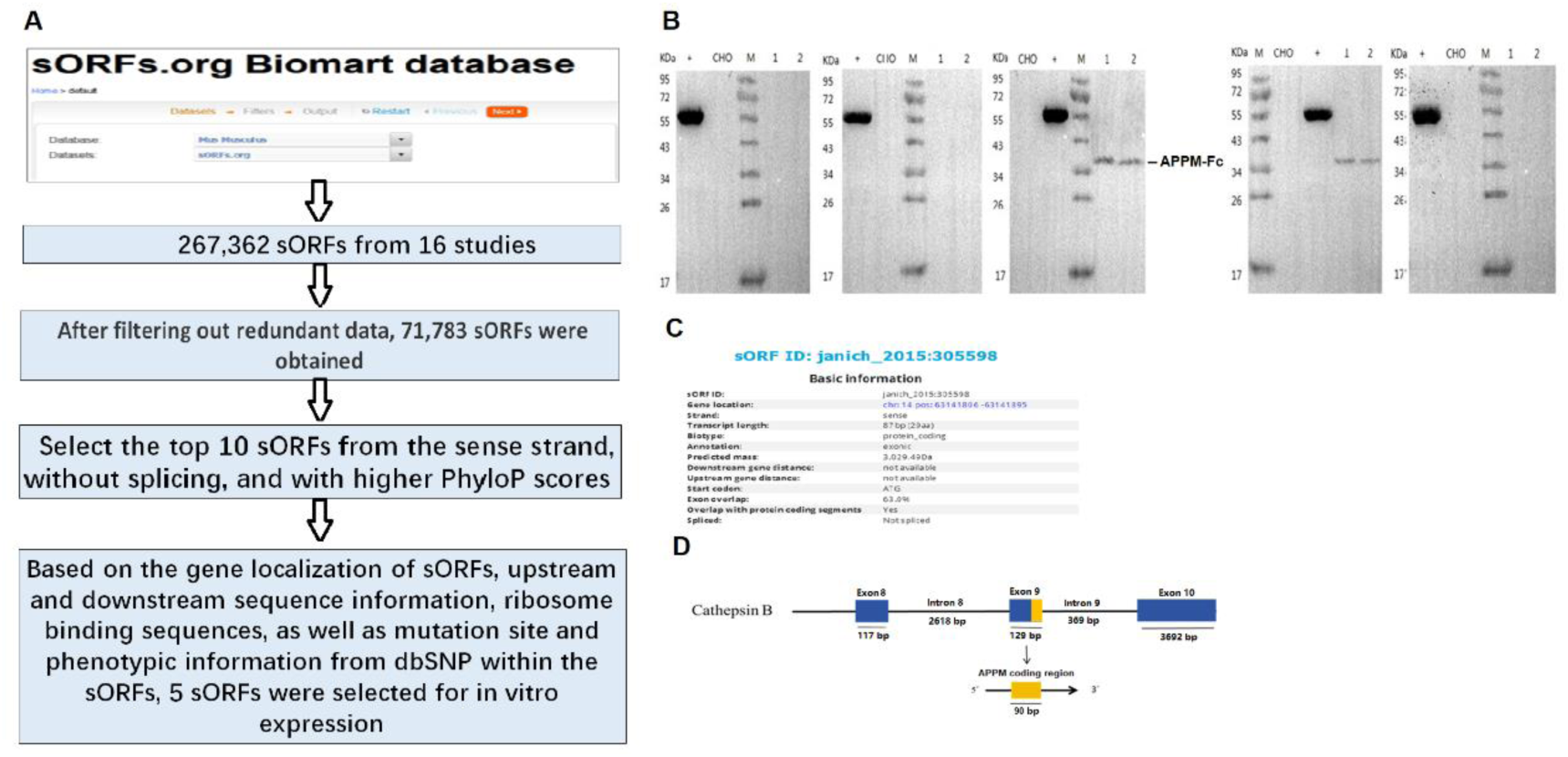
The APPM-encoded sORF screening from the sORFs.org database. **(A)** Screening workflow for candidate sORFs. **(B)** Screening for sORFs with coding capability through *in vitro* expression (M, Molecular weight marker. 1, ExpiCHO cell lysate. 2, Sample 1. 3, Sample 2. **+**, Positive control (hIgG heavy chain expressed in ExpiCHO). **(C)** The basic information of the APPM-encoded sORF. **(D)** The APPM-encoded sORF locates within the exonic and intronic regions of the CTSB protein-coding gene.

Collectively, these analyses identified a novel 90-nucleotide alternative open reading frame (alt-ORF) within the *CTSB* gene locus, which we named APPM, and confirmed its translation *in vitro*.

### Detection of micropeptide APPM translated from sORF in mice

To confirm endogenous production of APPM and obtain initial insight regarding its potential function, we validated its expression *in vivo* and investigated its tissue distribution and subcellular localization.

To this end, homozygous CRISPR/Cas9-mediated 3×Flag knock-in mice were generated to verify endogenous expression, (Fig. 2A, S1A, B). APPM-Flag fusion peptide and APPM were detected in the cardiac tissues of gene knock-in mice using anti-Flag antibodies and prepared anti-APPM monoclonal antibodies (Fig. S1E-I), respectively (Fig. S1C, D; Fig. 2B, C). Mass spectrometry analysis of mouse cardiac tissues further confirmed the existence of the endogenous micropeptide APPM (Fig. 2D). Notably, endogenous APPM exhibited an apparent molecular weight of approximately 27 kDa, substantially larger than that of the synthetic peptide (3.03 kDa). Bioinformatic prediction followed by experimental validation revealed O-glycosylation at Ser21, providing a mechanistic explanation for this size discrepancy (Fig. S1J). APPM was detected in several high-energy-demand organs, including the heart, liver, kidney, and brain (Fig. 1G, S1D). Based on tissue expression, N2a, HL-1 cell lines were selected for in *vitro* studies. Anti-APPM immunoblotting revealed low expression in N2a cells but high levels in HL-1 cells, mirroring tissue patterns (Fig. 2G). Subcellular fractionation of HL-1 cells showed APPM enrichment in the cytoplasm (Fig. 2F).

**Figure 2.**
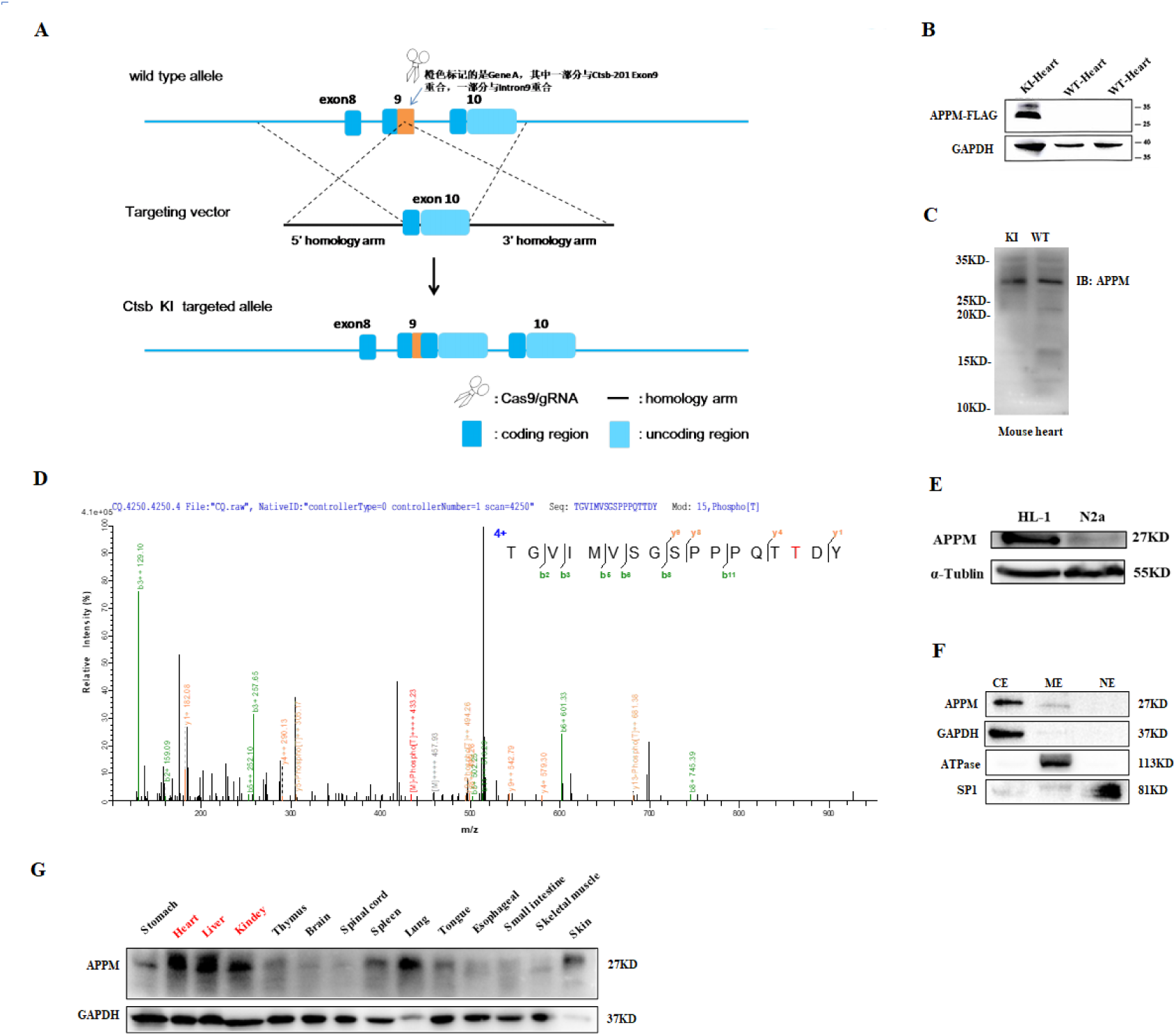
Identification and characterization of the endogenous micropeptide APPM. **(A)** The strategy for knocking in (KI) 3×Flag using CRISPR/Cas9 gene editing technology. **(B)** Detection of APPM-Flag fusion peptide expression in the hearts of flag KI mice using an anti-Flag antibody. **(C)** Detection of APPM in cardiac tissues from KI mice and wild-type mice using anti-APPM monoclonal antibodies. **(D)** LC-MS/MS spectrum showing the matched peaks of endogenous micropeptide APPM in mice heart. **(E)** Western blot analysis of APPM relative expression in HL-1 and N2a cell lines. **(F)** Cytoplasmic localization of APPM as identified from Subcellular fractionation (nuclear/NE, cytoplasmic/CE, and membrane/ME fractions). **(G)** Detection of APPM in various tissues of mice using an anti-APPM monoclonal antibody.

Together, these results conclusively verified the endogenous expression of APPM, revealed its specific enrichment in metabolically active tissues, and established its predominant localization within the cytoplasmic compartment.

### APPM directly interacts with OGDH

To elucidate APPM’s molecular function, co-immunoprecipitation (Co-IP) coupled with mass spectrometry was performed using cardiac tissue from *APPM–Flag* knock-in mice. Among 272 candidate interactors (Fig. S2A), Gene Ontology and KEGG pathway analyses highlighted strong enrichment for mitochondrial processes, particularly the TCA cycle and oxidative phosphorylation (Fig. S2B, C). Subsequent validation via reciprocal Co-IP identified OGDH, the E1 subunit of the OGDH complex, as a specific binding partner, with no interaction detected for the E2 (DLST) or E3 (DLD) subunits (Fig. 2A–J; Table S1). This interaction was conserved in wild-type tissues and various cell lines (Fig. S2B-D). The direct binding between synthetic APPM and recombinant OGDH was quantitatively confirmed by microscale thermophoresis (MST), revealing a dissociation constant (K_d_) of 1.44 × 10^−5^ M (Fig. S4). APPM overexpression had no effect on OGDH mRNA or protein levels. In contrast, OGDH knockdown promoted APPM expression, suggesting a potential negative feedback regulation in response to impaired energy production caused by OGDH deficiency (Fig. S2G, H). Confocal microscopy with MitoTracker Red confirmed mitochondrial colocalization of APPM (Fig. 3K). To further resolve its sub-mitochondrial localization, mitochondria were fractionated by osmotic shock followed by differential centrifugation, revealing enrichment of APPM within the mitochondrial inner membrane and matrix compartments (Fig. 3L, M). These results established OGDH as a direct and specific binding partner of APPM, confirmed their interaction within mitochondria, and localized APPM to the inner mitochondrial membrane and matrix, thereby linking it to core energy metabolism pathways.

**Figure 3.**
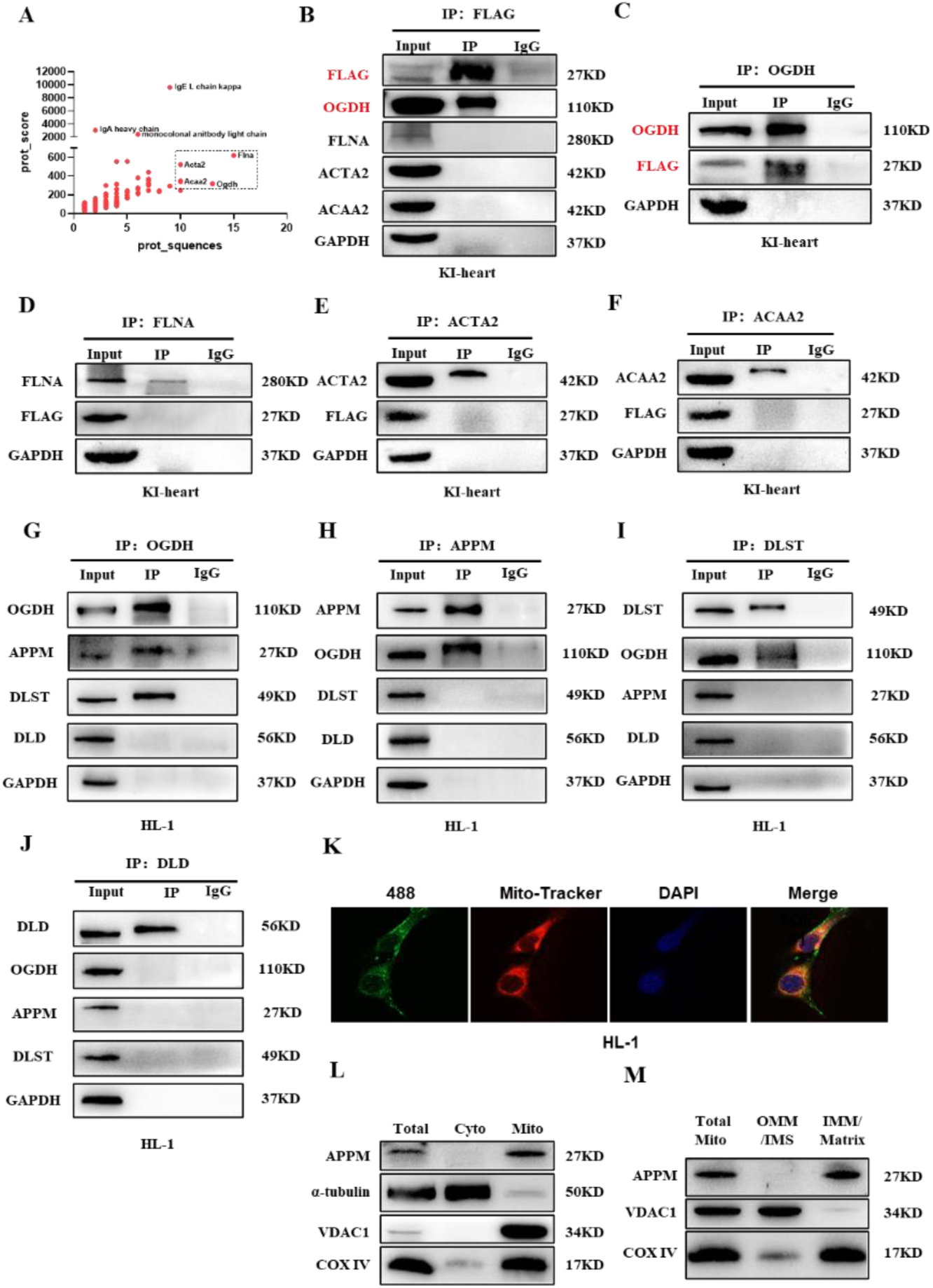
APPM directly interacts with OGDH and localizes to the mitochondrial matrix and inner membrane. **(A)** Screening of potential interacting proteins based on unique peptide counts and enrichment scores. **(B)** Co-immunoprecipitation (Co-IP) using anti-Flag antibody to enrich APPM-Flag fusion protein and its interacting partners from KI mice heart tissues. **(C-F)** Using antibodies against OGDH, FLNA, ACTA2, and ACAA2 for reverse co-IP, only anti-OGDH successfully enriched the APPM-Flag fusion protein from KI mouse heart tissues, while the other antibodies showed no enrichment. **(G-J)** Using antibodies against OGDH, APPM, DLST, and DLD for co-IP in HL-1 cells, APPM only interacts with E1 (OGDH) subunit of the OGDH complex. **(K)** Immunofluorescence (IF) and confocal microscopy reveal APPM colocalization with mitochondria (Mito-Tracker signal). **(L)** Mitochondrial isolation assay confirms APPM enrichment in the mitochondrial fraction. **(M)** Mitochondrial subfractionation shows APPM distribution near the inner mitochondrial membrane (IMM) and matrix. **Cyto**: Cytosol. **Mito**: Whole mitochondria. **VDAC1**: Mitochondrial loading control (outer membrane). **COX IV**: Mitochondrial marker (complex IV). **OMM/IMS**: Outer mitochondrial membrane and intermembrane space. **IMM/Matrix**: Inner mitochondrial membrane and matrix.

### Endogenous APPM regulates mitochondrial physiology, OGDHc activity, and ATP production

Since APPM is localized to the mitochondrial matrix and inner membrane and mitochondrial membrane potential, and morphology are closely linked to mitochondrial function, we examined the effect of APPM on mitochondrial morphology and bioenergetic status. Examination of mitochondrial ultrastructure via transmission electron microscopy (TEM) revealed that geonetic overexpression of APPM (Fig. S4A-G) caused mitochondrial swelling, consistent with elevated metabolic activity, whereas APPM knockdown resulted in smaller, presumably less active mitochondria (Fig. 4A). APPM overexpression selectively increased mitochondrial ATP levels (Fig. 4B, Table S2). whereas APPM knockdown led to a pronounced reduction in the mitochondrial membrane potential (ΔΨm), resulting in mitochondrial depolarization (Fig. 4C, D; Fig. S4H). These findings demonstrate that APPM is essential for preserving mitochondrial bioenergetic capacity and structural integrity, linking its molecular regulation of OGDHc to broader mitochondrial function.

**Figure 4.**
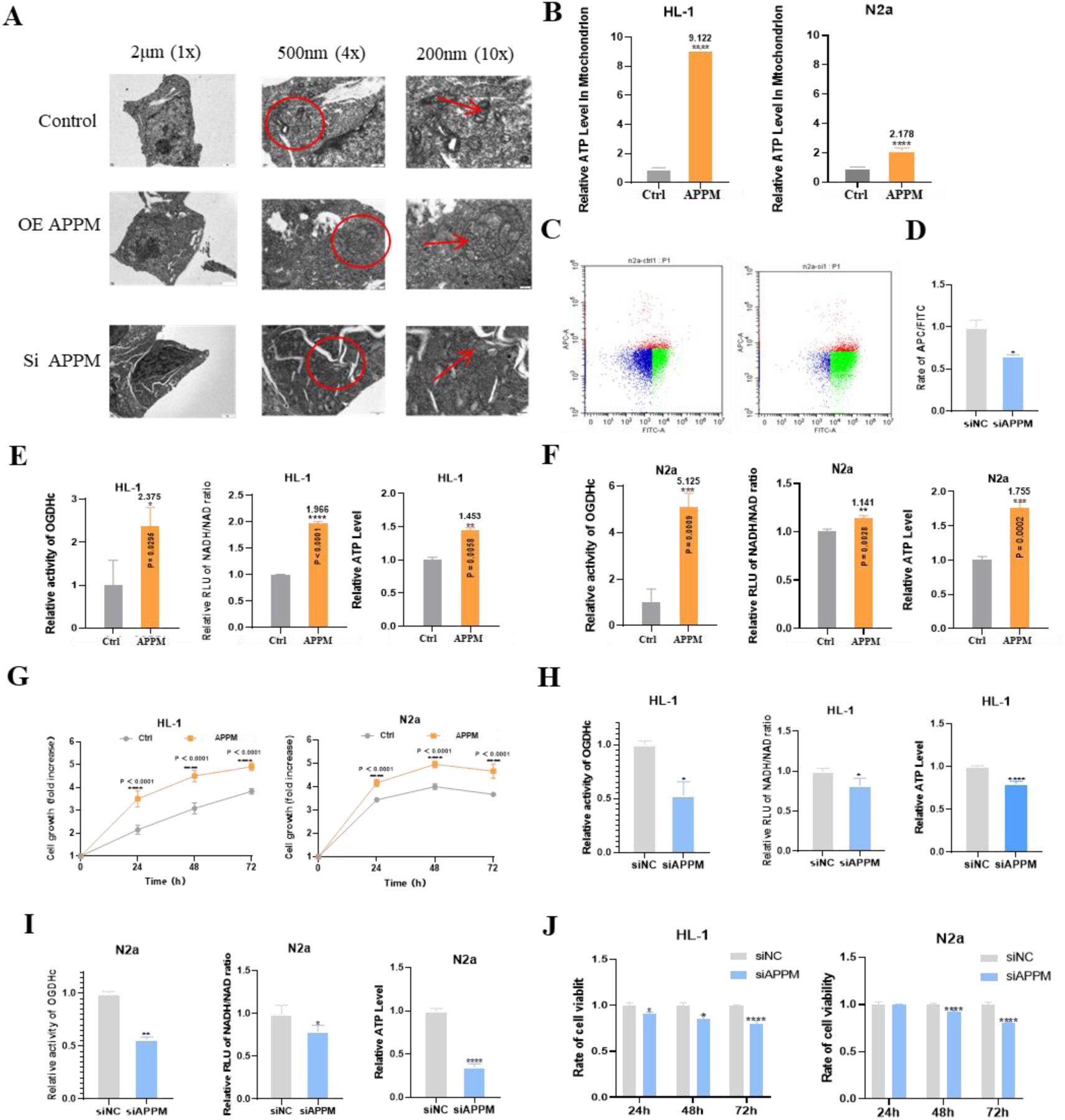
APPM regulates mitochondrial physiology, OGDHc activity, ATP production, and cell viability. **(A)** Transmission electron microscopy (TEM) revealing mitochondrial ultrastructural changes upon APPM overexpression (oeAPPM) or knockdown (siAPPM). **(B)** APPM overexpression increases relative ATP levels in HL-1 and N2a cells mitochondria. **(C)** Flow cytometry analysis of mitochondrial membrane potential in N2a cells after APPM **(D)** knockdown (siAPPM).Normal cells (red) and depolarized cells (green) were distinguished by red/green fluorescence intensity, reflecting mitochondrial depolarization. **(E-F)** APPM overexpression enhances OGDHc enzymatic activity, increases NADH/NAD⁺ ratio, and promotes ATP synthesis in HL-1 and N2a cells. **(G)** APPM overexpression stimulates cell proliferation in HL-1 and N2a cells. **(H-I)** Knockdown of APPM reduces OGDHc activity, NADH/NAD⁺ ratio, and ATP levels in HL-1 and N2a cells. **(J)** Knockdown of APPM inhibits cell proliferation in HL-1 and N2a cells.

Given the confirmed APPM-OGDH interaction, we next investigated its functional impact on OGDHc enzymatic activity and downstream energy metabolism in two high-energy-demand cell models: cardiomyocyte-derived HL-1 and neuron-like N2a cells. In both HL-1 and N2a cells, APPM overexpression significantly enhanced OGDHc enzymatic activity (1.1 to 5.1-fold increase; Fig. 4E, F, Table S2, S3). This boost in activity was accompanied by a corresponding 35-45% elevation in the NADH/NAD⁺ ratio (Fig. 4E, F, Table S2, S3), reflecting increased generation of reducing equivalents. Consequently, APPM augmentation led to a substantial rise in intracellular ATP concentration (1.1 to 2.1-fold; Fig. 4E, F, Table S2, S3) and a 60-75% enhancement in cell proliferation rates over 72 hours (Fig. 4G). Metabolite profiling further confirmed enhanced OGDHc flux: α-ketoglutarate (α-KG) levels decreased, whereas the product succinate accumulated in both cell lines, consistent with APPM-driven stimulation of TCA cycle turnover. (Fig. S5).

Conversely, RNA interference–mediated knockdown of APPM in HL-1 and N2a cells resulted in a marked reduction of OGDHc enzymatic activity (Fig. 4H, I), lowered the NADH/NAD⁺ ratio (Fig. 4H, I), and decreased intracellular ATP levels (Fig. 4H, I), accompanied by a significant decline in cell proliferation (Fig. 4J). Moreover, metabolite analysis revealed a delayed conversion of α-ketoglutarate (α-KG) to succinate (Fig. S5), consistent with impaired TCA cycle flux. Collectively, these results demonstrate that APPM is indispensable for maintaining OGDHc catalytic activity, and that its loss disrupts mitochondrial energy metabolism, leading to reduced ATP synthesis and cellular growth capacity.

Collectively, these results demonstrate that APPM is essential for maintaining mitochondrial membrane potential, ultrastructure, and bioenergetic capacity. It regulates OGDHc enzymatic activity, NADH/NAD⁺ ratios and ATP production, and modulates cell proliferation in high-energy-demand cells.

### Mapping the APPM-OGDH interaction interface

To delineate the structural basis of the APPM-OGDH interaction, we combined computational modeling and experimental validation. Molecular models of APPM and the OGDHc complex were constructed using AlphaFold3^40^ (Fig. S6A–C). Rosetta docking and molecular dynamics simulations predicted that APPM binds specifically to the OGDH monomer within the complex, with its N- and C-termini engaging the OxoGdeHyase_C and primary catalytic domains of OGDH, respectively (Fig. 5A, B). Notably, tyrosine at position 13 (Tyr13) was identified as critical for stabilizing the interaction.

**Figure 5.**
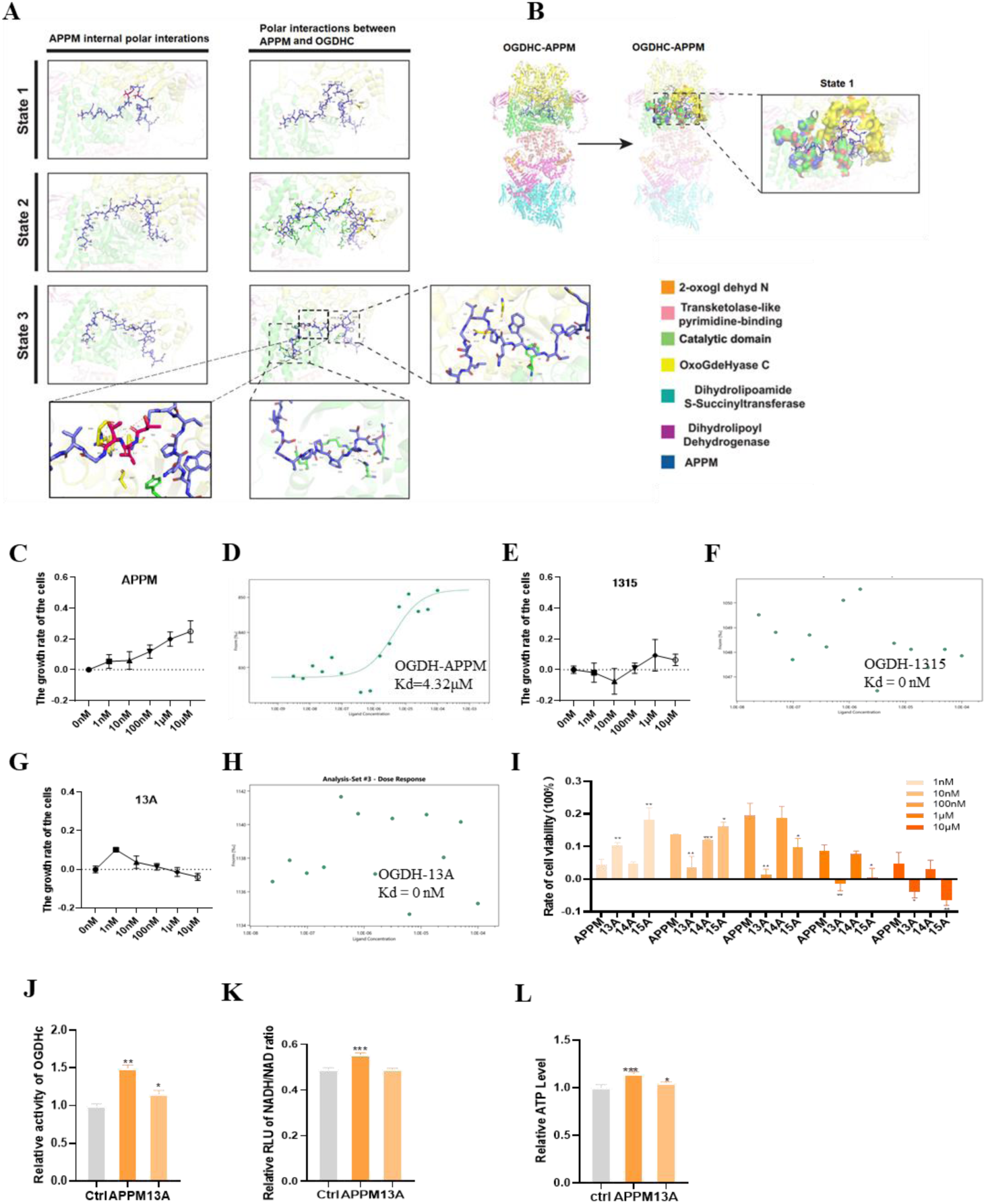
Identification of APPM’s active site and its binding interface with OGDH. **(A)** Detailed molecular interactions between OGDHc and APPM, highlighting the amino acids involved in the binding. **(B)** Binding configuration of the OGDHc complex with APPM. **(C)** Effects of wild-type APPM on cell proliferation at different concentrations. **(D)** MST binding assays validating interaction between OGDH and wild-type APPM peptides. **(E)** Effects of truncated peptide 13-15 on cell proliferation at different concentrations. **(F)** Truncated peptide 13-15 lose binding capacity to OGDH (MST assay validation). **(G)** Effects of 13-site alanine-substituted mutant peptide (13A) on cell proliferation at varying concentrations. **(H)** Mutant 13A peptide lose binding capacity to OGDH (MST assay validation). **(I)** Comparative analysis of cell viability under equivalent concentrations of different mutant peptides. **(J)** The mutant peptide (13A) showed weaker OGDHc enzyme activity upregulation than the original APPM peptide. **(K)** The mutant peptide (13A) did not affect intracellular NADH levels. **(L)** The mutant peptide (13A) slowed down intracellular ATP production. Data analyzed by GraphPad Prism 8.0 (unpaired t-test). *P<0.05, **P<0.01, ***P<0.001, ****P<0.0001. All experiments were repeated ≥3 times.

To experimentally validate these predictions, we performed systematic truncation and alanine-scanning mutagenesis of APPM. These analyses revealed that the peptide segment encompassing residues 13-15 was essential for function (Fig. 5C–F). Specifically, the Y13A mutation abolished OGDH binding (Fig. 5G, S6D, E), abrogated the proliferative effect of APPM (Fig. 5H), and impaired its ability to enhance OGDHc activity, ATP production, and α-KG to succinate conversion (Fig. 5J–L, S6F, G).

Collectively, these results demonstrate that Tyr13 serves as the structural linchpin mediating APPM-OGDH complex formation via salt bridge/hydrogen bond networks, thereby driving allosteric activation of TCA cycle flux and establishing a mechanistic foundation for metabolic rescue strategies.

### Synthesized APPM promotes ATP synthesis and stimulates cell proliferation in HL-1 and N2a cells

To systematically evaluate the physiological functions and therapeutic potential of chemically synthesized APPM, we synthesized the peptide using solid-phase peptide synthesis and performed comprehensive functional characterization (Fig. S3A, B). As shown in Fig. 6, the synthetic APPM peptide efficiently penetrated cells and selectively targeted mitochondria. Confocal microscopy revealed that FITC-labeled APPM readily crossed the plasma membrane and co-localized with MitoTracker, confirming mitochondrial accumulation (Fig. 6A, B). Functionally, exogenous APPM treatment dose-dependently enhanced the enzymatic activity of OGDHc in both HL-1 and N2a cells (Fig. 6C, D), increasing activity up to 1.415-fold and 1.451-fold, respectively, at 1000 nM. This was accompanied by a significant, dose-dependent increase in the cellular NADH/NAD⁺ ratio (Fig. 6E, F) and a corresponding promotion of ATP synthesis (Fig. 6G, H), with ATP levels rising up to 1.864-fold in HL-1 and 1.964-fold in N2a cells at the highest APPM concentration. Ultimately, these metabolic enhancements stimulated cell proliferation, as APPM treatment led to a substantial fold increase in the growth of both cell lines over a 60-hour period compared to controls (Fig. 6I, J). These findings demonstrate that the synthesized APPM mimics the activity of endogenous APPM, suggesting significant therapeutic potential for mitochondrial bioenergetic disorders.

**Figure 6.**
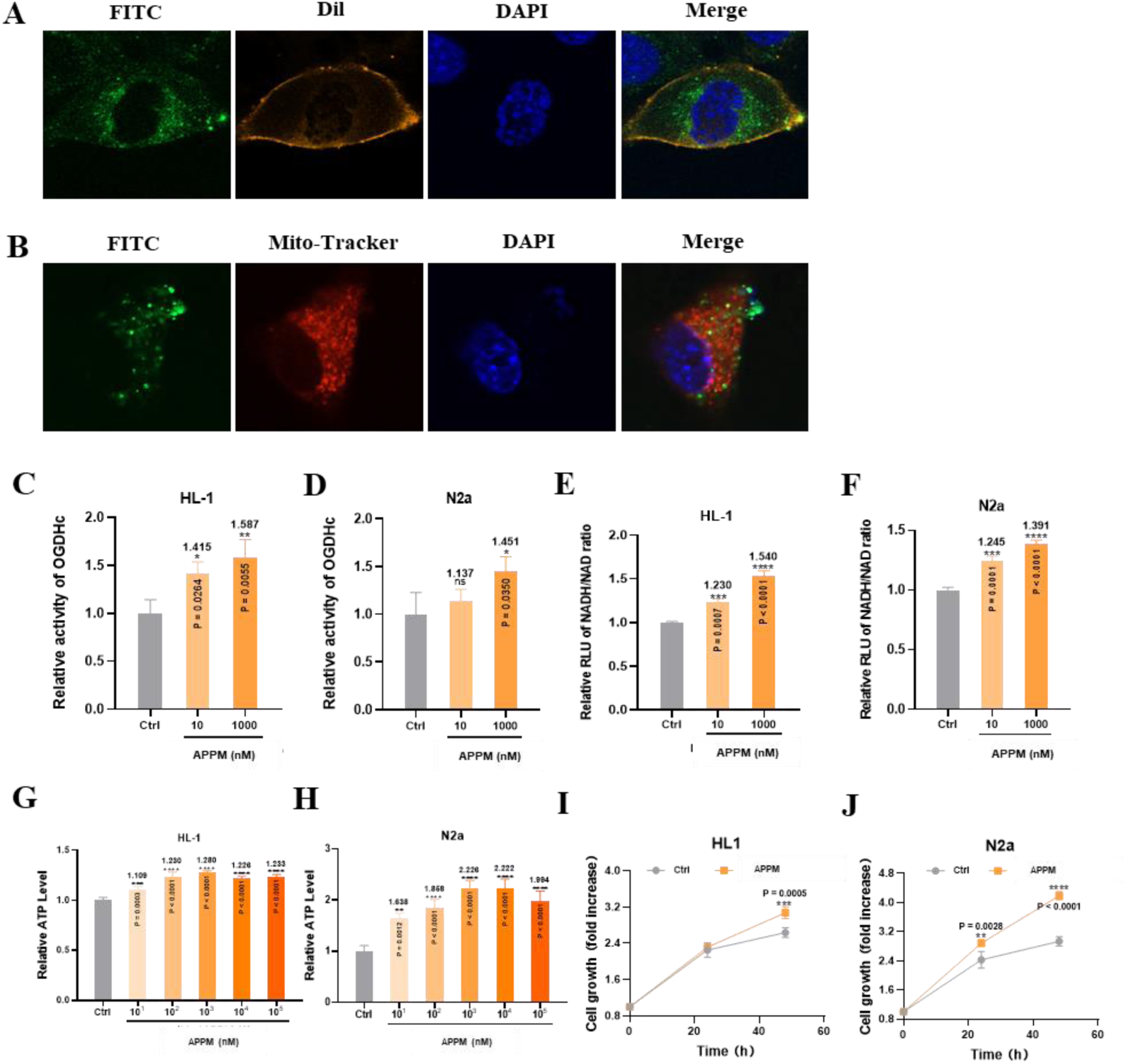
Synthesized APPM translocates across the plasma membrane and enter mitochondria to enhance OGDHc activity, increase NADH/NAD⁺ ratio, promote ATP synthesis, and stimulate cell proliferation. (A-B) Confocal microscopy demonstrated that FITC-labeled APPM can translocate across the plasma membrane and enter mitochondria. **(C-D)** Enhancement of OGDHc enzymatic activity by synthesized APPM in a dose-dependent manner. **(E-F)** Synthesized APPM increases NADH/NAD⁺ ratio in HL-1 and N2a cells in a dose-dependent manner. **(G-H)** Synthesized APPM promotes ATP synthesis in HL-1 and N2a cells in a dose-dependent manner. **(I-J)** HL-1 and N2a cells proliferation stimulation with Synthesized APPM (1000 nM).

### Therapeutic potential of APPM in heart failure

Heart failure is not only a mechanical failure of the pump function but is fundamentally characterized by a profound deficit in cellular energy metabolism. Correcting energy metabolism disorders has become an emerging research direction for treating heart failure. For example, the clinical efficacy of trimetazidine is related to its ability to inhibit fatty acid oxidation and promoting glucose oxidation^41^. APPM’s role in increasing ATP production by regulating OGDHc activity suggests its potential therapeutic value for heart failure. Therefore, we explored its therapeutic potential in heart failure. *In vivo* PET imaging of DOTA-labeled APPM showed rapid accumulation in the heart, liver, lungs, and kidneys, with the highest uptake in the kidneys (Fig. 7A, B). In normal mice, APPM injection (4.5 mg/kg) increased ATP levels in all tested organs within 4 hours, with a significant boost in brain OGDHc activity (Fig. 7C, D, Table S3).

**Figure 7.**
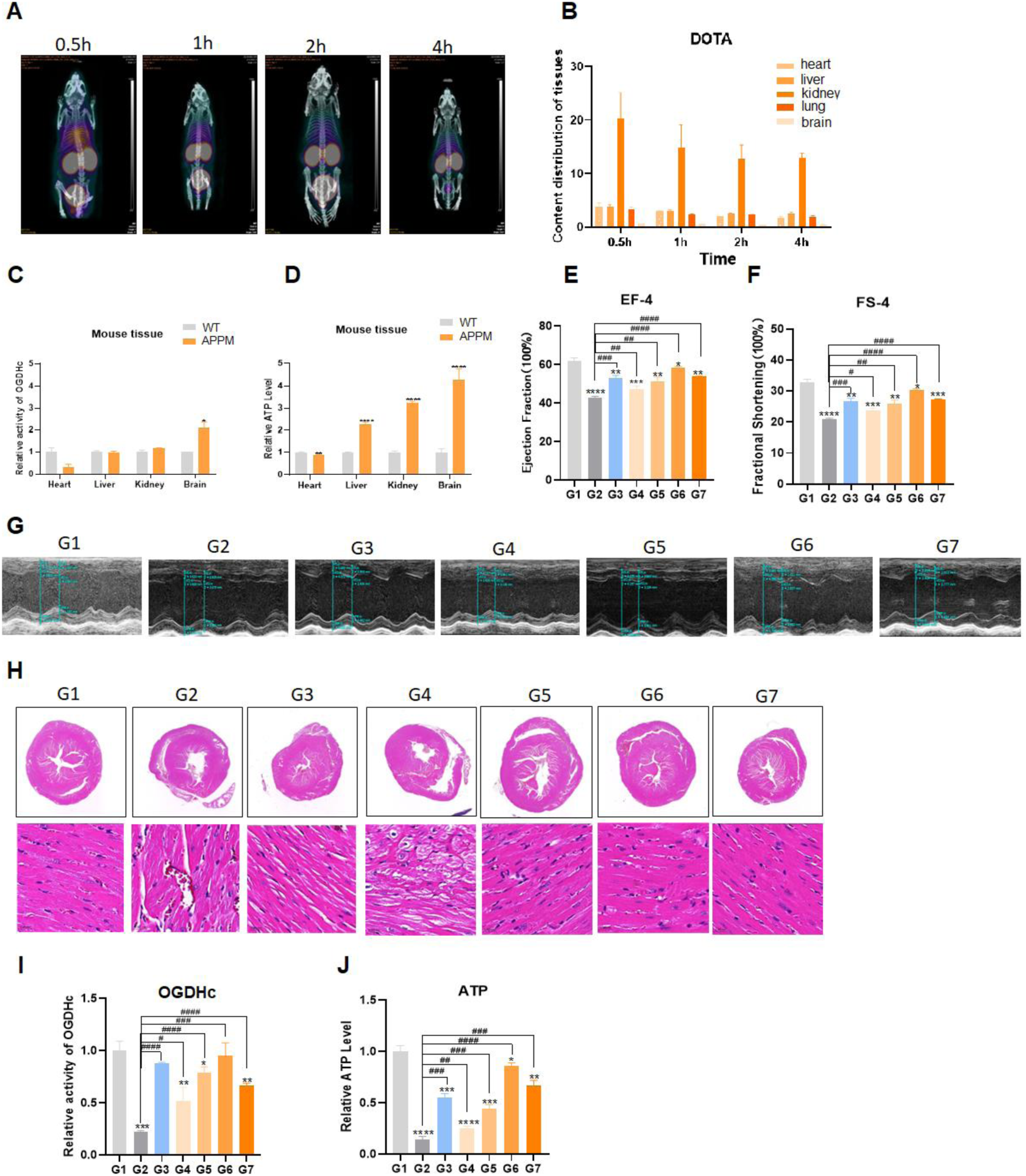
Synthetic APPM enhances ATP production in normal tissues and ameliorates isoproterenol-induced heart failure in mice. **(A-B)** Biodistribution analysis of DOTA-labeled APPM via PET imaging at 0.5, 1, 2, and 4 h post-injection. **(C-D)** Intravenous APPM (4.5 mg/kg) after 4 h increases OGDHc activity and ATP levels in heart, liver, kidney, and brain tissues. **(E-G)** Echocardiographic assessment after 1-month treatment: **(E-F)** APPM (4.5 mg/kg/day) stabilized ejection fraction (EF) and fractional shortening (FS) more effectively than positive control (trimetazidine, 5 mg/kg/day), though neither achieved complete recovery. **(G)** Representative images, **(H)** Histopathological improvements: APPM reduced cardiac dilation, inflammatory infiltration, and interstitial edema vs. heart failure model, outperforming trimetazidine. **(I-J)** APPM restored cardiac OGDHc activity and ATP production in failing hearts. Data were represented as mean ± SD (n=6). *vs control, #vs model group. Two-way ANOVA (GraphPad Prism 8.0). *P<0.05, **P<0.01, ***P<0.001, ****P<0.0001; #P<0.05, ##P<0.01, ###P<0.001, ####P<0.0001. All experiments were repeated ≥3 times.

We then employed an isoproterenol (ISO)-induced heart failure model. ISO administration successfully induced cardiac dysfunction, evidenced by structural damage, QRS prolongation, and ST-segment depression (Fig. S7A, B), and dynamic changes in endogenous APPM expression (Fig. S7C, D). Treatment with APPM for one week dose-dependently improved key cardiac functional parameters, including ejection fraction (EF) and fractional shortening (FS), with high-dose APPM (4.5 mg/kg/day) outperforming the reference drug trimetazidine (Fig. S7E–G, Table S4). After one month of treatment, APPM stabilized echocardiography parameters (EF, FS), with effects superior to trimetazidine (Fig. 7E–G, Table S4). Pathological features such as cardiac enlargement and inflammatory infiltration were significantly alleviated by APPM treatment (Fig. 7H). Crucially, APPM administration restored cardiac OGDHc activity and ATP production in the failing hearts (Fig. 7I, J), suggesting its therapeutic benefit is mediated through enhanced mitochondrial energy generation.

Excessive activation of β-adrenergic receptors in myocardial cells by isoproterenol triggers a series of pathological processes, including increased myocardial oxygen consumption, oxidative stress, and fibrosis. These changes lead to the gradual expansion of the extracellular matrix, impaired contractile function, and ultimately progression to heart failure. The micropeptide APPM binds to its interacting protein OGDH, activating the enzymatic activity of the OGDH complex. This action enhances the flux of the TCA cycle, promoting the generation of reducing equivalents and ATP in the mitochondria. Based on these metabolic regulatory mechanisms, APPM can regulate mitochondrial membrane potential, maintain mitochondrial morphology, improve cellular energy supply, thereby ameliorate heart failure progression.

## Discussion

This study identified and characterized APPM, a previously unrecognized mitochondrial micropeptide, derived from a non-canonical sORF within the CTSB gene. APPM was demonstrated to directly bind to the E1 subunit (OGDH) of OGHDc, thereby enhancing its enzymatic activity, increasing the NADH/NAD⁺ ratio, and promoting ATP production. To our knowledge, this represents the first identification of a novel protein component of this core enzyme complex in the TCA cycle since its elucidation, substantially expanding the current paradigm of mitochondrial energy metabolism.

Importantly, exogenous administration of synthetic APPM effectively improved mitochondrial dysfunction and cardiac contractile function in the mouse model of heart failure. Its efficacy was superior to that of the clinical control drug trimetazidine, and more importantly, it restored function to normal levels. This provides strong evidence for its potential application in treating bioenergetic metabolic dysfunction.

APPM is a novel component of the OGDHc. The TCA cycle is central to cellular energy metabolism, and its dysregulation is implicated in a wide range of pathological conditions, including neurodegenerative diseases, cardiovascular diseases, and metabolic syndrome^1–3^. OGDHc is one of the key rate-limiting enzymes in this cycle, and its activity is finely regulated by multiple mechanisms, including metabolite feedback, allosteric regulation, and post-translational modifications. This study reveals for the first time that APPM is an important component of OGDHc, acting through direct protein-protein interaction rather than indirect signaling pathways.

This mode of action distinguishes APPM from previously described mitochondrial micropeptides that indirectly affect energy metabolism (e.g., humanin, MOTS-c)^31,32^, APPM binds to OGDH through direct protein-protein interaction. Structural and mutational analyses further identified the tyrosine residue at position 13 of APPM as the critical site for maintaining its binding to OGDH and functional activation. The discovery of APPM not only adds a new dimension to TCA cycle regulation but also suggests that additional “hidden” components encoded by non-classical ORFs may remain undiscovered within even the most extensively studied metabolic pathways.

There may be a potential intrinsic connection and balanced relationship between APPM expression and CTSB gene expression. APPM in this study is encoded by an alt-ORF located within the CTSB gene locus, while CTSB itself encodes cathepsin B, a lysosomal protease involved in protein degradation and autophagy^42^. Interestingly, in the ISO-induced heart failure model, endogenous APPM expression exhibited dynamic regulation, suggesting transcriptional or translational control mechanisms distinct from those governing CTSB expression. Although APPM translation is independent of the main protein product of CTSB, their close genomic association may hint at synergistic or complementary regulatory relationships under certain physiological or pathological stress conditions.

Under energetic stress, upregulation of APPM may serve as a rapid adaptive mechanism to enhance ATP production, whereas CTSB plays a significant pathological role in promoting myocardial fibrosis, inflammation, and cardiomyocyte death during the onset and progression of heart failure. This implies a potential competitive process between APPM and the damage-promoting CTSB molecule, which might be a form of self-protection for the organism. Future research needs to deeply explore the characteristics of APPM’s own promoter/enhancer and whether its expression is difference between normal physiological and pathological states.

The effect of overexpressing APPM on mitochondrial ATP levels in cells, particularly the more than 9-fold increase in mitochondrial ATP levels upon APPM overexpression in HL-1 cells, far exceeds the fold increase in its enhancement of OGDHc enzymatic activity alone, indicating that APPM’s function likely extends beyond simple enzyme activation. We propose that APPM may amplify ATP synthesis by stabilizing the structure of the OGDHc complex, promoting its formation of supercomplexes with the mitochondrial inner membrane respiratory chain, or indirectly affecting the efficiency of other components of the electron transport chain.

These hypotheses align closely with emerging concepts in mitochondrial biology emphasizing the importance of metabolic enzyme organization, spatial compartmentalization, and supercomplex formation. This study found APPM distributed in high-energy-consuming organs and tissues such as the heart, liver, kidney, and brain. It may act as a “metabolic enhancement node,” optimizing the metabolic flux matching between the TCA cycle and oxidative phosphorylation to maximize energy output. This characteristic makes it an extremely valuable molecular tool for dissecting the precise regulatory mechanisms underlying explosive energy output in high-energy-demand cells (e.g., cardiomyocytes, neurons) under physiological and stress conditions.

The discovery of APPM has broad implications for mitochondrial research. First, as a micropeptide that directly targets and activates a core metabolic enzyme, APPM provides a controllable “molecular switch” for studying the dynamic regulation and physiological output of the TCA cycle. It can be used to precisely analyze the immediate and long-term impacts of metabolic flux changes on mitochondrial morphology, membrane potential, reactive oxygen species production, and cell fate. Second, APPM’s robust capacity to enhance ATP synthesis makes it an ideal probe for studying the limits of mitochondrial bioenergetics and exploring cellular energy supply strategies during regeneration, repair, or high-load conditions. Finally, given its nature as a small polypeptide and its proven cell-penetrating and mitochondrial-targeting abilities, APPM can serve as a carrier or lead molecule for developing novel mitochondrial-targeted delivery systems or designing superior synthetic micropeptides to modulate energy metabolism in specific disease states.

From a translational perspective, APPM represents a promising therapeutic candidate for diseases driven by mitochondrial energy insufficiency. Unlike conventional metabolic therapies that alter substrate utilization, APPM directly enhances TCA cycle flux, addressing cellular “energy starvation” at its core. Beyond heart failure, APPM or APPM-derived molecules may prove beneficial in neurodegenerative disorders, hereditary mitochondrial diseases, age-related metabolic decline, and acute energy crises such as ischemia–reperfusion injury (Fig. S9).

Future studies should focus on validating APPM function in human systems, optimizing its pharmacokinetic and delivery properties, assessing long-term safety, and exploring combination strategies with existing metabolic modulators. Collectively, our findings establish APPM as both a fundamental regulator of mitochondrial metabolism and a promising entry point for next-generation bioenergetic therapies.

## Methods

### Cell culture

N2a cells were cultured in Minimum Essential Medium (MEM) supplemented with 25 mM HEPES, 1% penicillin/streptomycin, 1% sodium pyruvate, and 10% fetal bovine serum (FBS). HL-1 cells were maintained in MEM containing 25 mM HEPES, 1% penicillin/streptomycin, and 10% FBS. All cell lines were maintained at 37°C in a humidified atmosphere containing 5% CO_2_.

### Identification and screening of small open reading frames (sORFs)

To identify candidate small open reading frames (sORFs), ribosome profiling data was obtained from the sORFs.org database, comprising 267,362 entries from 16 studies in mice. The dataset was filtered using the following stringent criteria: removal of redundant entries (sORFs reported in multiple studies), presence of ATG as the start codon, origin from the sense strand, and absence of alternative splicing in the transcript. From the filtered dataset, the top 10 sORFs were selected based on PhyloP conservation scores. These candidates were further analyzed for chromosomal localization, genomic context (upstream and downstream regions), and predicted ribosome binding sequences. Five sORFs were selected for in vitro expression validation, leading to the identification of two sORFs with detectable translation products. Among these, an 87-nucleotide sORF located within the exonic and intronic regions of the CTSB protein-coding gene was selected as the primary subject of this study.

### *In vitro* validation of sORF coding potential

Expression vectors were constructed based on the pcDNA3.1 backbone. These vectors included the 5’ UTR of APPM, the candidate sORF sequence (without its native stop codon), an Fc fragment coding sequence (677 bp), and a termination codon (TAA). Recombinant plasmids were transfected into CHO cells, which were maintained for 48-72h under standard culture conditions. Immunoblotting was performed using anti-Fc antibodies to validate translation by detecting the expected fusion protein. This system allowed for sensitive detection of sORF-encoded peptides through signal amplification mediated by the Fc fragment. Positive signals confirmed the translation capability of the candidate sORFs.

### Generation of Flag KI mice

A donor vector containing a 3×Flag coding sequence was designed using In-Fusion cloning technology. The insertion site was strategically targeted immediately upstream of the APPM stop codon to preserve endogenous gene regulation. Cas9 mRNA and guide RNA (gRNA) targeting the APPM locus were synthesized through *in vitro* transcription. The Cas9 mRNA, gRNA, and donor vector were co-injected into fertilized C57BL/6J mouse oocytes. The injected embryos were subsequently implanted into pseudopregnant females to produce F0 founder mice. F0 mice were screened for successful Flag knock-in using PCR and sequencing techniques. Flag-positive F0 mice were crossed with wild-type C57BL/6J mice to generate F1 heterozygotes. These F1 heterozygotes were then intercrossed to produce F2 homozygous mice. (Note: Flag knock-in heterozygous mice were generated by Shanghai Model Organisms Center.)

### Micropeptide synthesis

The synthetic micropeptide (purity >95%) was synthesized by GL Biochem (Shanghai) Ltd. using standard Fmoc solid-phase peptide synthesis (SPPS) methodology. The micropeptide was stored at-80°C desiccated until use.

### Immunoprecipitation (IP) for APPM enrichment

Total protein was extracted from N2a cells using a commercial extraction kit. The lysate was adjusted to a concentration of 1 mg/mL with protein lysis buffer. For immunoprecipitation, 500 μL of the diluted protein solution was incubated with 5–10 μL of target-specific antibody (at a dilution of 1:50–1:100). An IgG antibody was used as a negative control. The mixture was rotated gently at 4 °C for 16 h. Subsequently, 30–40 μL of Protein A/G agarose beads were added, and incubation continued at 4°C for 3 h with rotation. The samples were then centrifuged at 6,000 × g for 2 min at 4°C, and the supernatant was discarded. The beads were washed five times with 1 mL of TBS buffer using gentle inversion, followed by centrifugation at 6,000 × g for 1 min at 4°C. After washing, the beads were resuspended in 30–40 μL of RIPA lysis buffer and heated at 100°C for 10 min in a metal bath. The samples were centrifuged at maximum speed in a microcentrifuge for 3 min, and the supernatant was carefully transferred to a new tube, leaving the agarose beads behind. The supernatant could be stored at −80°C for further analysis. For glycosidase digestion of enriched APPM, 30 μg of the immunoprecipitated glycoprotein was denatured and then mixed with appropriate reagents for subsequent processing.

### O-Glycosylation analysis of APPM

The immunoprecipitated glycoprotein (30 μg) was denatured in 10 μL of solution containing 1 μL of 10× Glycoprotein Denaturing Buffer and ddH₂O, by incubation at 75°C for 10 min. Subsequently, 2 μL of 10× GlycoBuffer 2, 2 μL of 10% NP-40, and 1–4 μL of O-Glycosidase (optimized per manufacturer’s protocol) were added, and the total volume was adjusted to 20 μL with ddH₂O. The mixture was incubated at 37°C for 1–4 h. The reaction was terminated, and the samples were analyzed by SDS-PAGE or Western blotting to detect O-glycosylation alterations.

### LC-MS/MS analysis

Heart tissue of 3×Flag mice were lysed with RIPA buffer for 30 min and then mixed with the SDS sample loading buffer. The proteins were separated by SDS-PAGE. The protein bands were excised, decolorized, reduced and digested in the gel. The samples were dissolved in 0.1% TFA and 2% acetonitrile, and the resulting peptides were analyzed by QExactive mass spectrometer (Thermo Fisher Scientific) and high-performance liquid chromatograph (Dionex Ultimate 3000 RSLCnano). The linear gradients are (A) 0.1% formic acid, (B) 0.1% formic acid, 80% acetonitrile, and the flow rate is 300 nL/min. The Q Exactive mass spectrometer was operated in data-dependent mode. Full scans were acquired at a resolution of 70,000 over the m/z range 350-1800. The top 20 most intense precursor ions were selected for fragmentation via higher-energy collisional dissociation (HCD) and identified by tandem mass spectrometry (MS/MS).

### Lentiviral packaging and transfection

1×10^7^ 293T cells were cultured in 15 cm cell culture dishes in 10 mL of medium (containing 1% penicillin/streptomycin and 10% bovine serum). The lentiviral vector containing cDNA sequence (pLVX-mCMV-ZsGreen1-Puro, addgene) was co-transfected with psPAX2 (addgene) and PMD2. After 72 h, the supernatant carrying lentivirus was collected and filtered through a 0.45 μm filter screen to remove cell debris. The supernatant was then concentrated using the Fast-Trap Lentivirus Purification and Concentration Kit (Millipore) according to the manufacturer’s protocol. Two days after infection, puromycin with a final concentration of 2 µg/mL was added to screen for stable transfected cell lines.

### Gene knockdown

A mixture of 100 μL Opti-MEM and 12.5 μL of either negative control or target gene siRNA was prepared and mixed thoroughly by vortexing. Separately, 100 μL of Opti-MEM was combi ned with 5 μL of liposomal transfection reagent by gentle shaking (vortexing was avoided). The two mixtures were then combined and incubated at room temperature for 20 minutes to form transfection complexes. Meanwhile, the cell culture medium was replaced with 1.3 mL of fresh complete medium. Subsequently, 200 μL of the transfection complex was added to the cells. The cultures were incubated at 37°C with 5% CO₂ for 24 hours, after which the medium w as changed. Gene expression analysis was performed 48 to 72 hours post-transfection. All procedures were conducted using RNase-free consumables to prevent contamination. siRNA sequenc es (synthesized by Genepharma): Ctsbmus21 (N2a): Forward: 5′GCAGCCAACUCUUGGAACCT T3′; Reverse: 5′GGUUCCAAGAGUUGGCUGCTT3′; APPM (HL1): Forward: 5′CCCUCCUCAA ACUACAUAA3′; Reverse: 5′UUAUGUAGUUUGAGGAGGGGG3′.

### Quantitative PCR (qPCR)

Total RNA was isolated using TRIzol (Invitrogen) and reverse transcribed with the High-Capacity cDNA Reverse Transcription Kit (Takara). qPCR was performed using SYBR Green Master Mix (Takara) on a QuantStudio 3 system (Applied Biosystems). Relative expression (fold change) was calculated via the 2^(ΔΔCt) method, normalized to GAPDH for both cellular and tissue lysates.

### Co-immunoprecipitation (Co-IP)

Heart lysates from 3×Flag APPM mice were prepared in RIPA buffer (Thermo) with protease inhibitors. After centrifugation (12,000 g, 15 min, 4°C), supernatants were immunoprecipitated overnight at 4°C with anti-Flag antibody (14793, Cell Signaling Technology). Protein complexes were captured using Protein A/G agarose beads (9863, Cell Signaling Technology), washed 3× with RIPA buffer, and analyzed by SDS-PAGE followed by LC-MS/MS or western blot.

### Western blotting

Cells/tissues were lysed in RIPA buffer containing protease/phosphatase inhibitors (Thermo). Proteins (20 μg) were separated by SDS-PAGE, transferred to PVDF membranes (Millipore), blocked with 5% milk, and probed overnight at 4°C with primary antibodies: Flag (Abcam), OGDH (Cell Signaling Technology), FLNA/ACTA2/ACAA2 (Santa Cruz), ATPase (Proteintech), SP1/VDAC1/COX IV (Wanleibio). HRP conjugated secondary antibodies and ECL were used for detection. Band intensity was quantified using ImageJ.

### Microscale thermophoresis (MST)

OGDH (1 μM) was labeled with fluorescent dye (Ruixin Biotech) for 90 min at RT. Serial twofold dilutions of APPM peptide (200 μM initial in PBS) were prepared. Binding assays were performed on a Monolith NT.115 (NanoTemper) and analyzed with Nano Temper Analysis software.

### APPM monoclonal antibody production

Mice were immunized with the conjugated antigen peptide. Sera were screened by ELISA, and a titer >8,000 was required to proceed with hybridoma fusion. Hybridoma fusion was performed, and cells were subcloned until 100% antigen-positive clones were obtained (Performed by Abmart, Shanghai).

### Subcellular fractionation

A total of 1×10⁶ cells were washed once with pre-cooled PBS. Discard the supernatant, add 300 μL of CEB (Thermo), incubate at 4°C for 10 min, centrifuge at 500 g at 4°C for 5 min, and take the supernatant as the cytoplasmic extract. After the precipitate was washed once with PBS, 300 μL MEB (Thermo) was added, incubated at 4 °C for 10 min, centrifuged at 3000 g at 4°C for 5 min, and the supernatant was the cell membrane extract. Add 150 μL of NEB (Thermo), incubate at 4 °C for 30 min, centrifuge at 5000 g at 4°C for 5 min, and the supernatant is the nuclear extract.

### Mitochondrial isolation

A total of 2×10⁷ cells were resuspended in 800 μL of Reagent A (from a mitochondrial isolation kit) and incubated on ice for 2 min. Then, 10 μL of Reagent B was added, and the suspension was homogenized with 80 strokes. Subsequently, 800 μL of Reagent C was added. The homogenate was centrifuged sequentially at 700 ×g for 10 min, 3,000 ×g for 15 min, and 12,000 ×g for 5 min to pellet the mitochondria.

### OGDH activity & ATP measurement

OGDH activity was measured using a commercial kit (Mlbio). ATP levels were quantified with an ATP assay kit (Beyotime). NADH/NAD⁺ ratios were determined using a Promega kit.

### Mitochondrial membrane potential (JC10 Assay)

N2a cells were stained with JC10 (Yeasen Biotech) and analyzed by fluorescence microscopy (490/525 nm and 540/595 nm) and flow cytometry (FL1/FL2 channels). Mitochondrial morphology was assessed by TEM (JEM1400).

### Transmission electron microscopy (TEM) for mitochondrial morphology analysis

Approximately 10 million cells were centrifuged at 300 g for 5 min at 4°C in a 1.5 mL microcentrifuge tube. The cell pellet was carefully resuspended in a freshly prepared solution of 2.5% glutaraldehyde in 0.1 M sodium cacodylate buffer (pH 7.4). Fixation was allowed to proceed for 12 h at 4 ° C with continuous gentle agitation. Subsequently, the sample was centrifuged at 3,000 g for 5 min to consolidate the fixed cells into a cohesive pellet. For storage and transport, samples were maintained in the fixative at 4°C. During transport, insulated containers with wet ice were used, and an adequate volume of fixative was ensured to prevent desiccation. The samples were rinsed three times with 0.1 M sodium cacodylate buffer (pH 7.4). Post-fixation was performed with 1% osmium tetroxide in 0.1 M sodium cacodylate buffer for 2 h at 4°C. The samples were then rinsed three times with distilled water. Dehydration was carried out through a graded ethanol series (30%, 50%, 70%, 90%, 100%) at 4°C, allowing 20 min per step. The samples were then transitioned to propylene oxide twice, for 15 min each. Infiltration with Epon 812 resin mixtures was performed as follows: a 1:1 mixture of resin to propylene oxide for 2 h at room temperature, followed by a 2:1 mixture overnight at room temperature, and finally with pure resin for 8 h at room temperature. Polymerization was conducted at 60°C for 48 h.

Sections of 70 nm thickness were prepared using a diamond knife ultramicrotome and subsequently collected on 200 mesh copper grids. The sections were stained with 2% uranyl acetate in 50% ethanol for 8 min, followed by counterstaining with Reynolds’ lead citrate for an additional 8 min. Examination of the sections was conducted using a JEM-1400 transmission electron microscope operating at an accelerating voltage of 80 kV. Digital images were captured at magnifications ranging from 10,000× to 50,000×. For each sample, a minimum of ten representative fields were acquired. Magnification calibration was performed using carbon grating replicas.

### ISO induced disease model

Six-week-old male C57BL/6J mice (n=35) were acclimated for 7 days before ISO injection (85 mg/kg on Day 1; 340 mg/kg on Day 2, s.c.). Mortality was approximaely 50%. Cardiac function was assessed by ECG on Day 10.

### Echocardiography

Anesthetized mice (1.5-2% isoflurane) were imaged using a highresolution ultrasound system (30-50 MHz probe). Bmode (long/short axis) and Mmode (EF, FS) measurements were acquired, with pulsewave Doppler for hemodynamic parameters (E/A ratio, CO). Core temperature was maintained at 37°C; triplicate measurements were averaged.

### Electrocardiography (ECG)

ECGs were recorded under light anesthesia (1-2% isoflurane) using subcutaneous needle electrodes (Lead II configuration) and a PowerLab system (ADInstruments; sampling rate ≥1 kHz). HR, PR, QRS, and QT intervals were analyzed from ≥5 stable cycles using LabChart.

### Molecular Structure Modeling and Molecular Docking

We utilized the online version of AlphaFold3 server for molecular structure modeling. Rosetta software was employed for flexible molecular docking. Molecular dynamics (MD) simulations were performed using GROMACS 2022 with the GROMOS54A7 force field. The simulation was set at a temperature of 310 K and pressure of 101 kPa in a neutral environment. Temperature and pressure equilibration were conducted for 1 ns with a time step of 2 fs.

### Quantification and statistical analysis

Data for quantification analyses are represented as mean ± SEM. Significance of difference was examined by Student’s t test (*P < 0.05, **P < 0.01, ***P < 0.001) or one-way ANOVA followed by Fisher’s LSD multiple-comparison test. Different letters at top of columns indicate significant differences at P < 0.05. The statistical analyses for all experiments were performed with the GraphPad Prism 8 software. The exact value of n, what n represents and the statistical details of each experiment are described in the figure legends or tables.

## Data availability

All data are available in the main text and the supplemental information or at public databases. This study does not report original code.

Any additional information required to reanalyze the data reported in this study is available from the lead contact upon request.

## Acknowledgments

This work was supported by the University-Industry Collaboration Program (No. 8210040001, 2020-019) from Nanjing Anji Biotechnology Co., Ltd. Also thanks to the National Natural Science Foundation (No. 82373039), 2024 Annual Research Project on Multi-Target Natural Medicines (State Key Laboratory Project) (SKLNMZZ2024JS08), Team Development for Peptide-Based Pharmaceutical Research and Development (1132230001A002), Peptide Drug Research and Development Track (3342400029). The authors gratefully acknowledge Dr. Mengwei Li for her insightful guidance and support in this study.

## Author contributions

H.X. designed and supervised the study; X.Y., and H.X. discovered micropeptide APPM; X.Y., and Q.C. performed the screening and validation of APPM *in vitro* and *invivo*; L.L. carried out loss-of-function studies, investigated the mechanistic basis of micropeptide activity, and identified critical functional residues. H.Zh., and G.Zh. performed experiments including subcellular localization of the micropeptide, interaction protein analysis, overexpression assays, and functional characterization of chemically synthesized peptides. G.Sh. conducted radiolabeling experiments to determine the *in vivo* distribution of the micropeptide. Y.W. performed molecular structure modeling and molecular docking analyses. H.X., X.Y., W.Q., and J.H. wrote and revised the paper; H.X. acquired funding. All authors have read and agreed to the published version of the manuscript.

