## Supplemental tables and figures for "A micropeptide component of the OGDH complex modulates ATP production in the TCA cycle"

### Supplemental data

#### Supplemental table

Table S1 Top 30 APPM potential interacting proteins ranked by unique peptide count

| Gene Name | UniProt | Description | Mass | Score | Unique Sequences |
|---|---|---|---|---|---|
| Flna | B7FAU9 | Filamin, alpha | 283177 | 621 | 15 |
| Ogdh | Q60597 | 2-oxoglutarate dehydrogenase, mitochondrial | 117572 | 318 | 13 |
| Acta2 | Q3U122 | ACTA protein | 42367 | 522 | 10 |
| Acaa2 | Q3UKH3 | 3-ketoacyl-CoA thiolase, mitochondrial | 42259 | 349 | 10 |
| Hsp90ab1 | Q71LX8 | Heat shock protein 84b | 83571 | 341 | 10 |
| Pkm | P52480 | Pyruvate kinase | 58378 | 248 | 10 |
| Tubb4b | P68372 | Tubulin beta-4B chain | 50255 | 293 | 9 |
| Tubb5 | P99024 | Tubulin beta-5 chain | 50095 | 243 | 8 |
| Pdia3 | P27773 | Protein disulfide-isomerase | 57099 | 235 | 8 |
| Hbbt1 | A8DUP7 | Beta-globin | 15930 | 442 | 7 |
| Actb | Q3UBP6 | ACTB protein | 42084 | 368 | 7 |
| Ckm | A2RTA0 | Creatine kinase | 43246 | 349 | 7 |
| Immt | Q3TEY5 | MICOS complex subunit | 75924 | 308 | 7 |
| Hspd1 | P63038 | 60 kDa heat shock protein, mitochondrial | 61088 | 302 | 7 |
| Krt8 | P11679 | Keratin, type II cytoskeletal 8 | 54531 | 247 | 7 |
| Tuba1c | Q3TIZ0 | Tubulin alpha chain | 50592 | 381 | 6 |
| Tuba4a | A0A0A0MQA5 | Tubulin alpha chain (Fragment) | 53612 | 375 | 6 |
| Tpm2 | Q6PJ18 | Tpm2 protein | 32995 | 274 | 6 |
| Atp1a1 | Q8VDN2 | Sodium/potassium-transporting ATPase subunit alpha-1 | 114221 | 266 | 6 |
| Hsp90b1 | Q3UAD6 | Histidine kinase | 92703 | 221 | 6 |
| Ckb | Q04447 | Creatine kinase B-type | 42971 | 212 | 6 |
| Pdia6 | Q3TJL8 | Protein disulfide-isomerase A6 | 49039 | 244 | 5 |
| Got1 | P05201 | Aspartate aminotransferase, cytoplasmic | 46504 | 226 | 5 |
| Hspa5 | Q3TI47 | 78 kDa glucose-regulated protein | 72415 | 194 | 5 |
| Clu | Q549A5 | Clusterin | 52250 | 168 | 5 |
| P4hb | Q3TF72 | Protein disulfide-isomerase | 57452 | 166 | 5 |
| Aldoa | A6ZI44 | Fructose-bisphosphate aldolase | 45548 | 313 | 4 |
| EG433182 | Q5FW97 | Phosphor-pyruvate hydratase | 47453 | 259 | 4 |
| Eno3 | Q4FK59 | Phosphor-pyruvate hydratase | 47310 | 214 | 4 |
| Anxa5 | P48036 | Annexin A5 | 35787 | 175 | 4 |

Table S2 The different biological effects of endogenous overexpression of the micropeptide APPM

| Cell | Model Processing | Fold Change | Enhanced OGDHc Enzyme Activity | Promoted Reduction | Promoted Cellular ATP Synthesis | Promoted Mitochondrial ATP Synthesis |
|---|---|---|---|---|---|---|
| N2a | Endogenous Overexpression | 2.32 | 5.125 | 1.141 | 1.755 | 2.178 |
| HL-1 | Endogenous Overexpression | 1.97 | 2.375 | 1.966 | 1.453 | 9.122 |

Table S3 The different biological effects of exogenous administration of the chemically synthesized peptide APPM

| Cell | Model Processing | Concentration (nM) | Enhanced OGDHc Enzyme Activity | Promoted Reduction | Promoted Cellular ATP Synthesis |
|---|---|---|---|---|---|
| N2a | Exogenous administration | 10 | 1.137 | 1.245 | 1.638 |
| | Exogenous administration | 1000 | 1.451 | 1.391 | 2.226 |
| HL-1 | Exogenous administration | 10 | 1.415 | 1.230 | 1.109 |
| | Exogenous administration | 1000 | 1.581 | 1.540 | 1.280 |

Table S4 Summary table of key indicator changes during the treatment of heart failure with micropeptide APPM (n=6)

| Time (days) | | G1 Control | G2 Model | G3 Positive drug (Trimetazidine 5mg/kg/day) | G4 APPM 0.5mg/kg /day (i.v.) | G5 APPM 1.5mg/kg /day (i.v.) | G6 APPM 4.5mg/kg /day (i.v.) | G7 APPM 4.5mg/kg /day (s.c.) |
|---|---|---|---|---|---|---|---|---|
| 3 | EF | 66.8±2.83 | 42.5±7.43 | 53.6±2.25 | 47.3±4.75 | 56.4±3.42 | 60.3±4.07 | 57.7±5.90 |
| | FS | 36.0±2.22 | 20.4±4.05 | 26.9±1.53 | 23.1±2.84 | 28.7±2.24 | 31.4±2.62 | 29.6±3.95 |
| 7 | EF | 65.9±2.98 | 49.4±4.63 | 49.5±5.97 | 53.2±3.21 | 54.8±4.05 | 61.1±2.43 | 65.9±12.3 |
| | FS | 35.5±2.49 | 24.5±2.81 | 24.5±3.53 | 26.6±1.67 | 27.7±2.66 | 32.0±1.92 | 35.8±9.36 |
| 30 | EF | 62.0±1.61 | 42.7±0.868 | 52.9±1.57 | 47.5±1.54 | 51.4±2.15 | 58.2±0.759 | 53.9±0.216 |
| | FS | 32.8±1.01 | 20.8±0.472 | 26.8±0.906 | 23.6±0.996 | 25.8±1.27 | 30.3±0.404 | 27.4±0.0565 |

### Supplemental figures

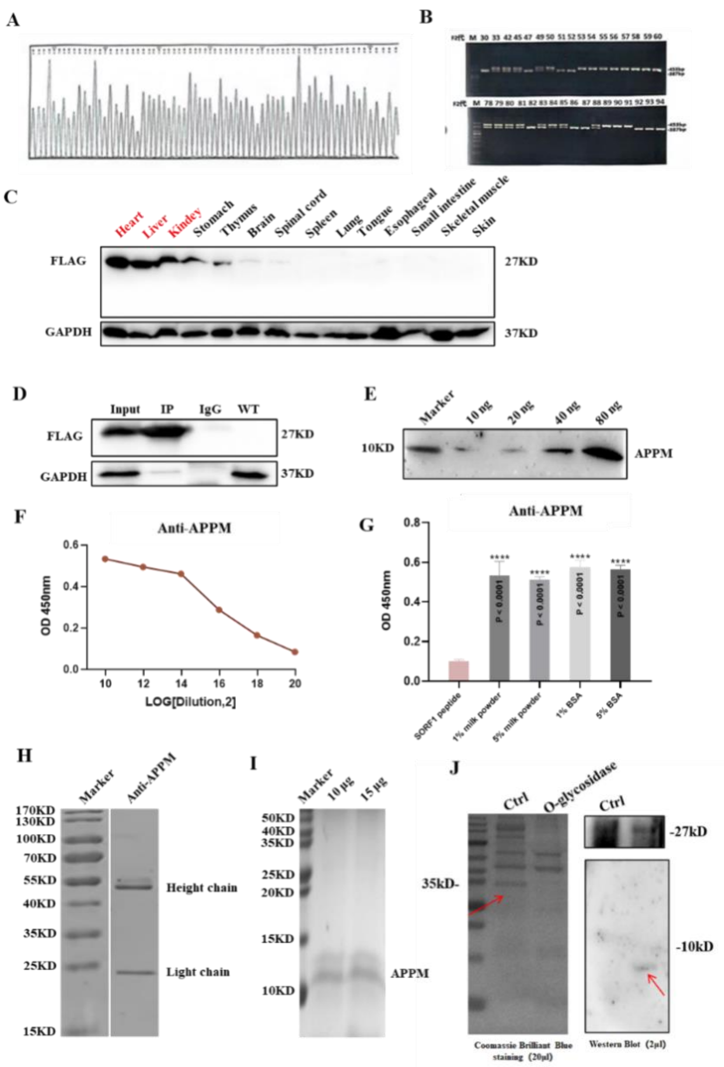


**Figure S1. Construction and validation of the flag knock-in mouse model, preparation of anti-APPM monoclonal antibody.**

(A) Sanger sequencing confirming the successful integration of the 3×Flag sequence.

(B) PCR genotyping of F2 generation homozygous mice carrying the 3×Flag knock-in.

(C) Detection of APPM-Flag fusion peptide expression in various tissues of flag KI mice using an anti-Flag antibody.

(D) Co-immunoprecipitation assay demonstrating that the APPM-encoding gene is co-translated with the 3×Flag tag, producing a Flag-tagged micropeptide.

(E) Western blot analysis of anti-APPM monoclonal antibody reactivity with immunogenic peptide at different concentrations.

(F) Anti-APPM monoclonal antibody titer determination.

(G) Anti-APPM monoclonal antibody specificity assessment by Western blot.

(H) Anti-APPM monoclonal antibody purity analysis by SDS-PAGE.

(I) Purity verification of the immunogenic peptide by SDS-PAGE.

(J) Validation of endogenous APPM O-glycosylation following O-glycosidase treatment.

All data represent mean ± SEM from three independent experiments. Statistical analysis was performed using one-way ANOVA in GraphPad Prism 8.0.

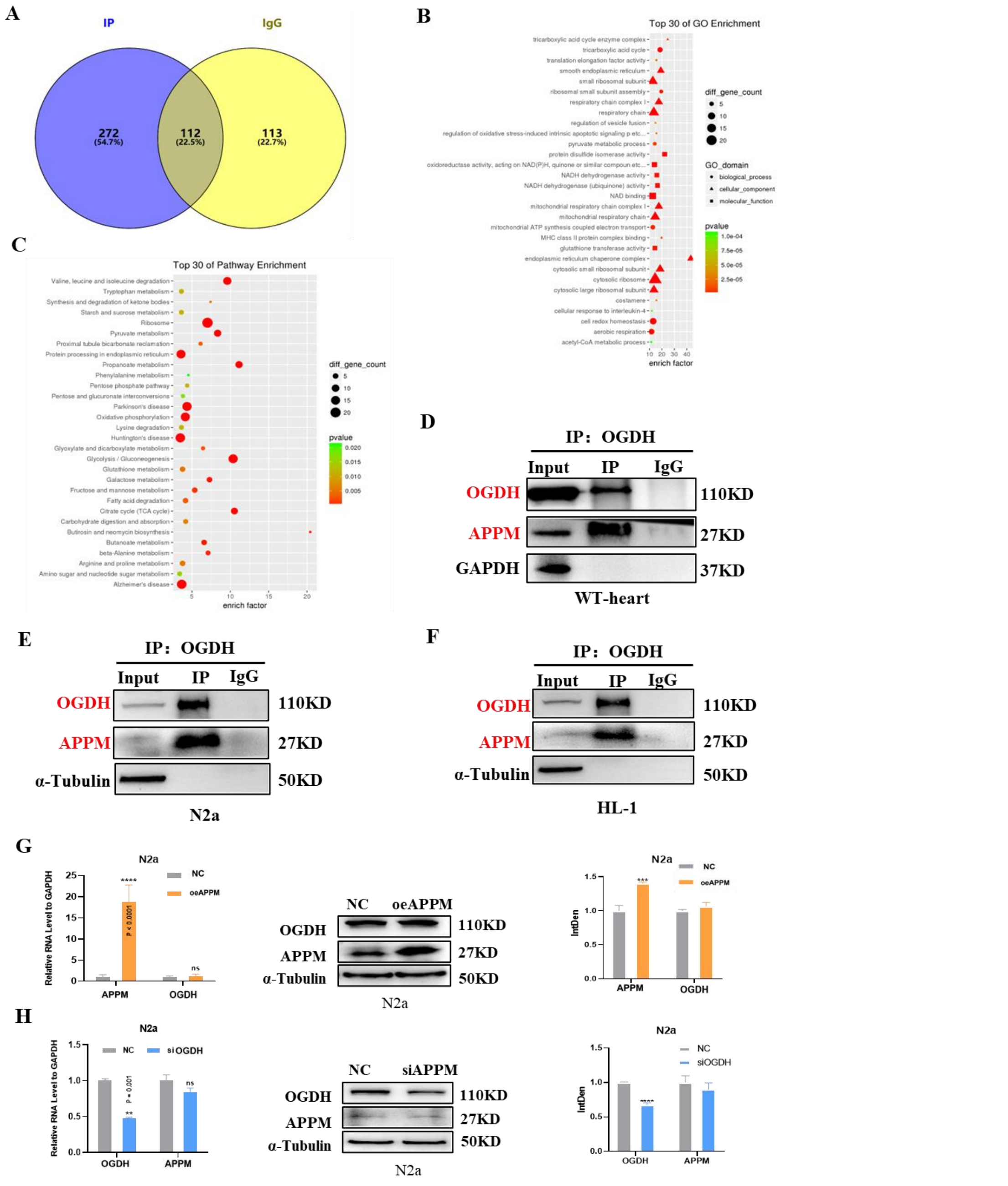


**Fig S2. Analysis of proteins interacting with APPM.**

**(A)** Venn diagram showing the overlap of APPM-interacting proteins identified by LC-MS/MS.

**(B)** Gene Ontology (GO) enrichment analysis of the APPM-interacting protein set.

**(C)** Kyoto Encyclopedia of Genes and Genomes (KEGG) pathway enrichment analysis of the APPM-interacting proteins.

**(D)** Reverse Co-IP using anti-OGDH antibody to enrich micropeptide APPM in wild-type mouse heart tissue.

**(E)** Reverse Co-IP using anti-OGDH antibody to enrich micropeptide APPM in N2a cells.

**(F)** Reverse Co-IP using anti-OGDH antibody to enrich micropeptide APPM in HL-1 cells.

**(G)** mRNA and protein levels of OGDH in N2a cells overexpressing APPM.

**(H)** mRNA and protein expression levels of APPM in N2a cells after OGDH knockdown.

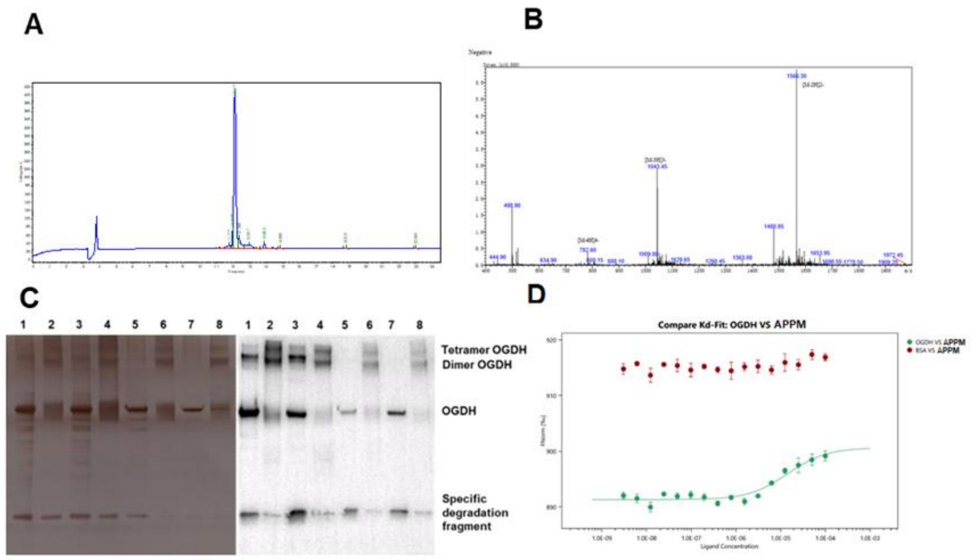


**Figure S3. Microscale thermophoresis (MST) analysis reveals direct binding between micropeptide APPM and protein OGDH related with figure 2.**

**(A)** HPLC quantification of chemically synthesized APPM peptide purity.

**(B)** Mass spectrometry (MS) verification of the molecular weight of synthetic APPM peptide.

**(C)** Expression and purification of OGDH protein from suspension-cultured HEK293F cells.

**(D)** MST-based binding assay demonstrating direct interaction between APPM and OGDH.

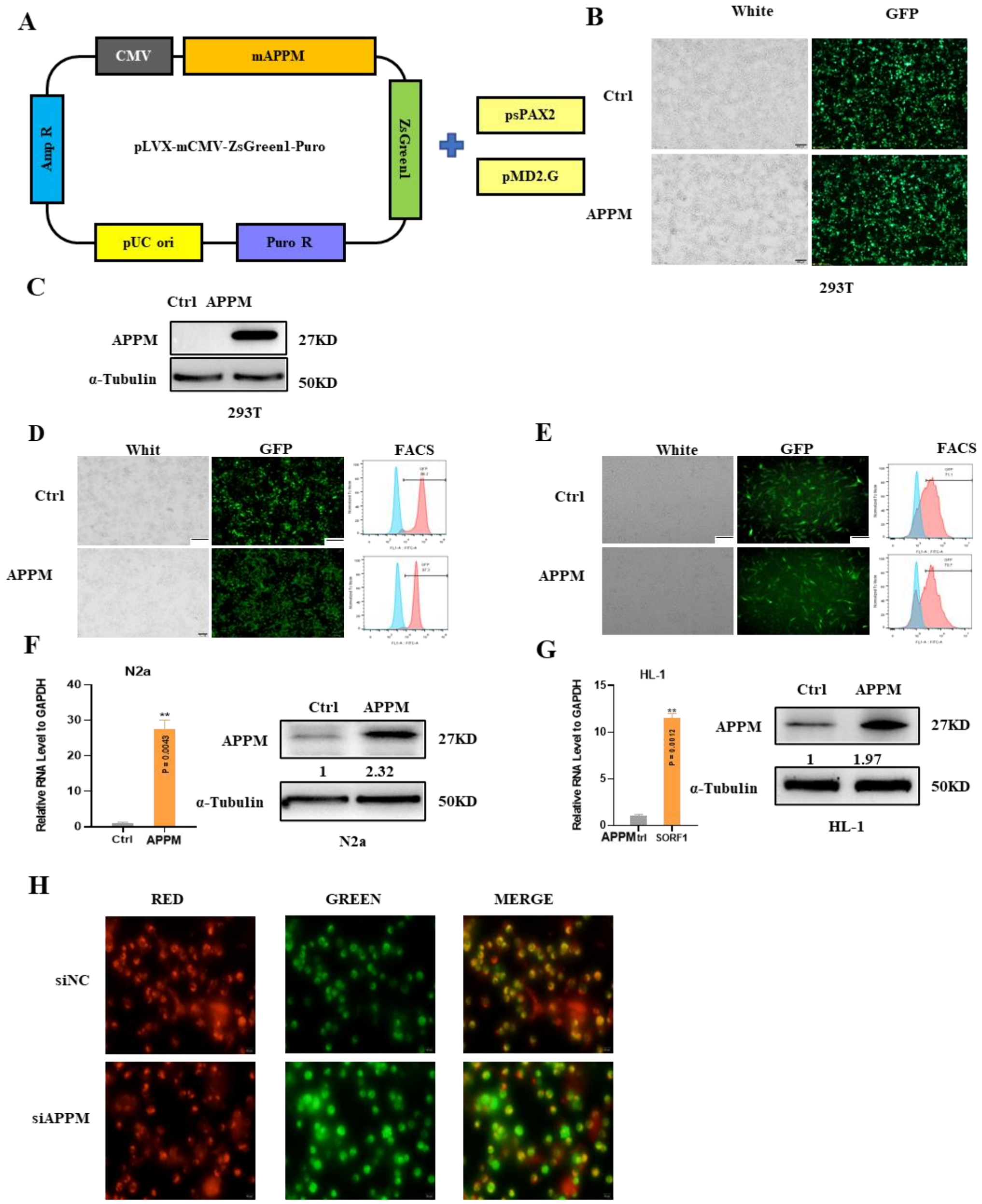


**Figure S4. Lentiviral expression and packaging in 293T cells, .**

**(A)** Schematic diagram of the lentiviral expression vector and packaging plasmids.

**(B)** GFP expression observed in 293T cells 48 h post-transfection.

**(C)** Western blot analysis of APPM protein expression in 293T cell lysates 72 h post-transfection.

**(D-E)** GFP expression efficiency in N2a and HL-1 cells assessed by **(D)** fluorescence microscopy and **(E)** flow cytometry.

**(F-G)** Validation of APPM overexpression in N2a and HL-1 cells by Western blot.

**(H)** Fluorescence microscopy showing differential red/green fluorescence intensity changes of mitocondria in Na2 cells.

All data represent mean ± SEM from three independent experiments. Statistical significance: $^{**}P < 0.01$.

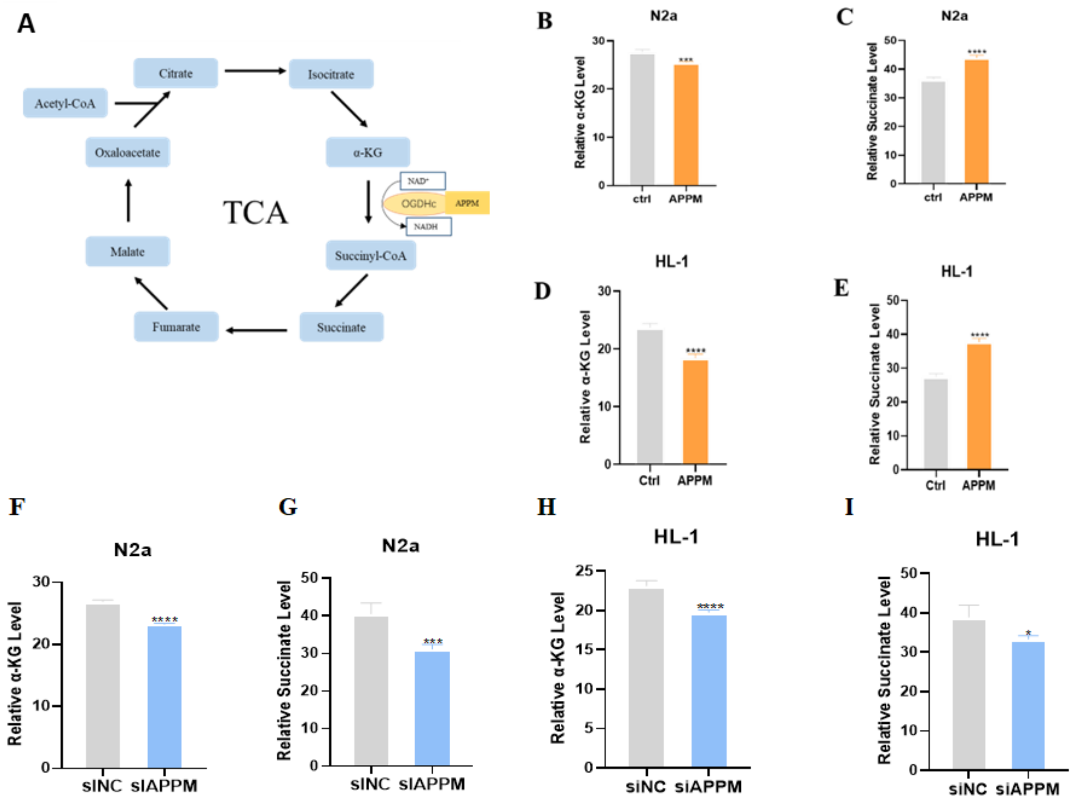


**Figure S5. APPM modulates α-KG and succinate production in HL-1 and N2a cell lines.**

**(A)** TCA cycle and electron transport chain; (B, D) Overexpression of APPM downregulates α-KG in HL-1 and N2a cell lines; (C, E) Overexpression of APPM upregulates succinate in HL-1 and N2a cell lines. (F, H) Knockdown of APPM inhibits intracellular α-KG production; (G, I) Knockdown of APPM suppresses intracellular succinate production. GraphPad Prism 8.0, T-test, *$P < 0.05$, ***$P < 0.001$, ****$P < 0.0001$.

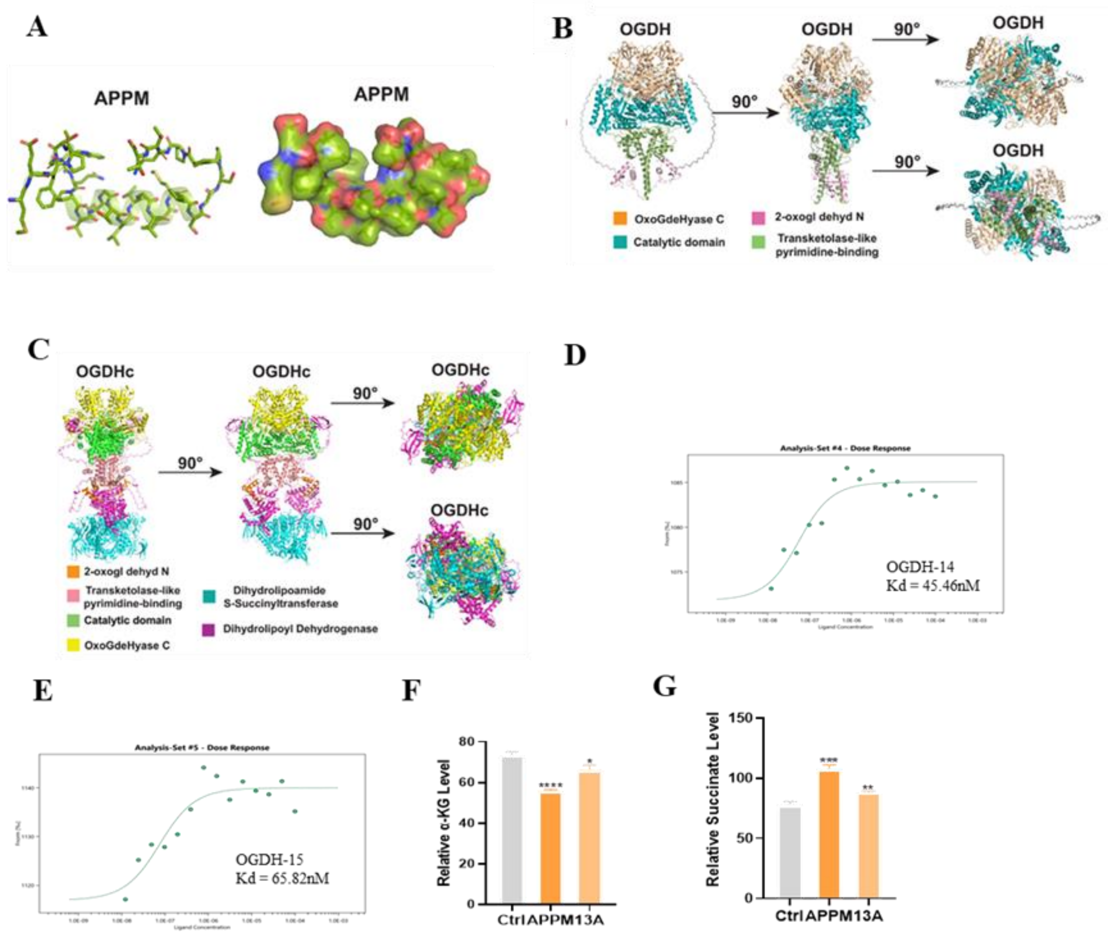


**Figure S6. Molecular Structural Characterization**

**(A)** Tertiary structure of the APPM polypeptide.

**(B)** Molecular tertiary structure of OGDH, with precise functional domain delineation.

**(C)** Quaternary structure of the OGDHc complex.

**(D-E)** MST binding assays validating interaction between OGDH and APPM mutant peptides.

**(F-G)** The micropeptide APPM accelerates the conversion of α-KG to succinate, while the 13th amino acid mutant peptide slows down the conversion of α-KG to succinate.

Data analyzed by two-way ANOVA using GraphPad Prism 8.0 (**P*<0.05, ***P*<0.01, ****P*<0.001, , *****P*<0.0001). All experiments were performed with at least three biological replicates.

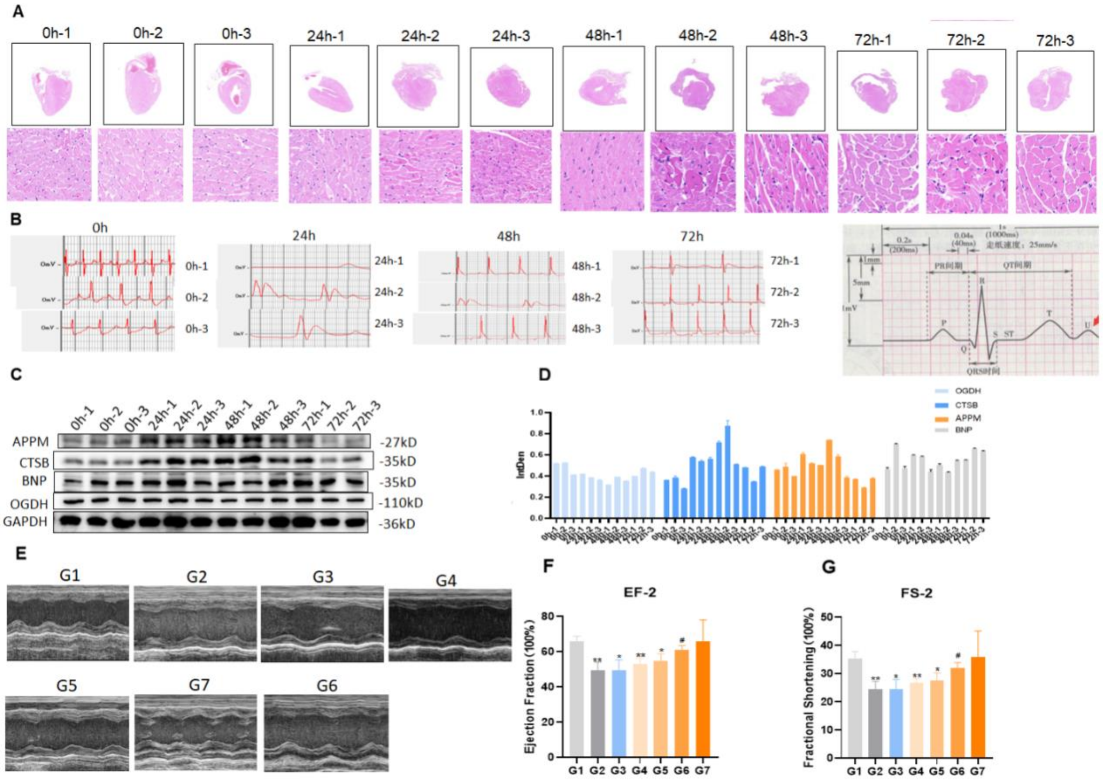


**Figure S7. Establishment of isoproterenol-induced heart failure mouse model and therapeutic evaluation of APPM peptide related with figure 6.**

**(A)** H&E staining showing cardiac tissue morphology changes at different time points (0h, 24h, 48h, 72h) post-isoproterenol induction.

**(B)** ECG recordings from heart failure model mice at four time points (0h, 24h, 48h, 72h) compared with normal ECG template.

**(C)** Western blot analysis of APPM and related protein expression during heart failure progression.

**(D)** Quantitative densitometry analysis of Western blot results.

**(E)** Representative echocardiograms after one week of peptide treatment.

**(F-G)** Echocardiographic parameters showing dose-dependent improvement in **(F)** ejection fraction (EF, normal range 55%-75%) and **(G)** fractional shortening (FS, normal range 25%-45%) following peptide treatment, with superior efficacy compared to positive control drug (trimetazidine, 5mg/kg/d).

Data are presented as mean ± SEM (n=6). Statistical analysis was performed using GraphPad Prism 8.0 (unpaired t-test). *P<0.05, **P<0.01 vs control group; #P<0.05 vs model group.

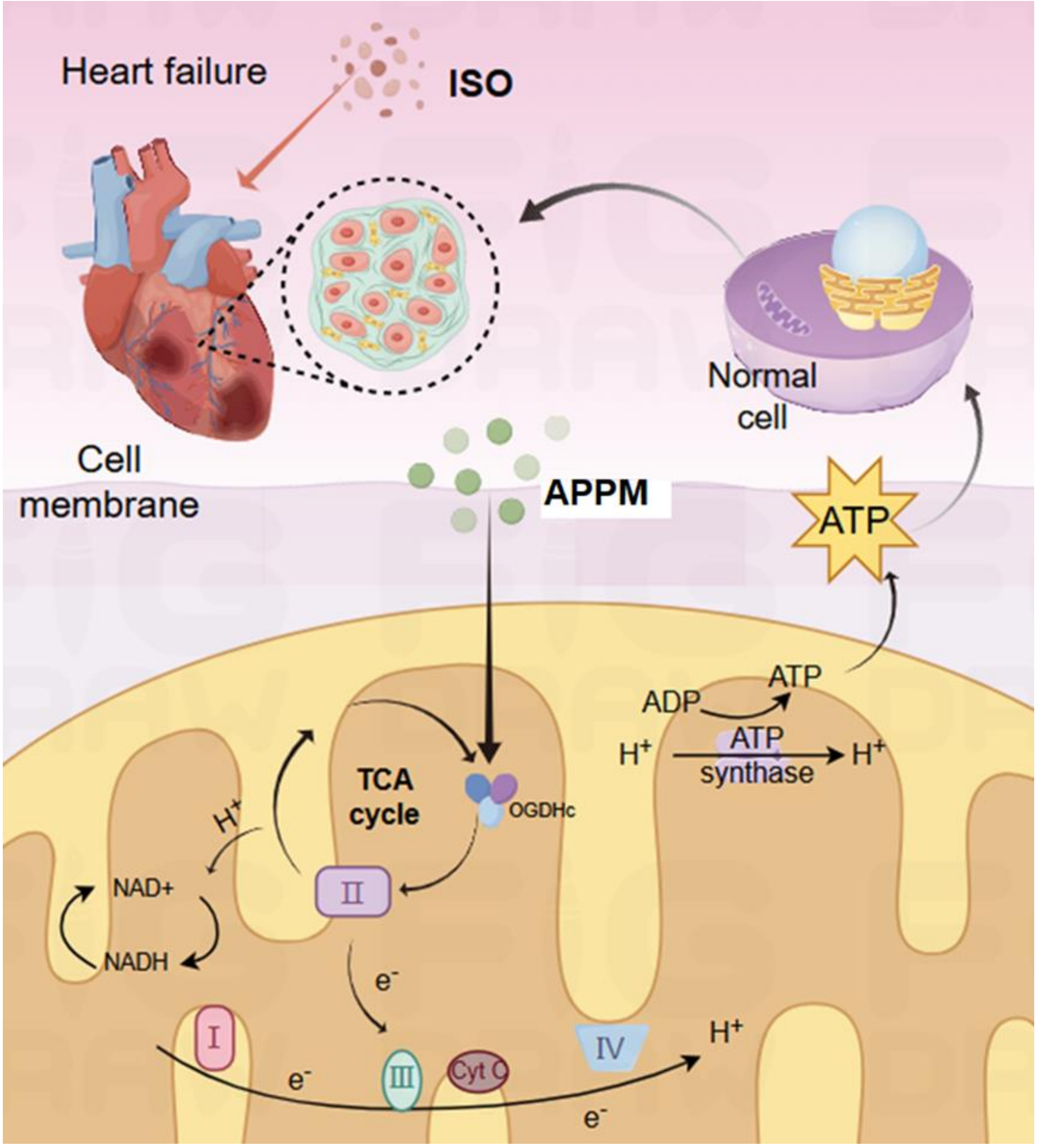


**Figure S8. Proposed mechanism of micropeptide APPM in alleviating heart failure.**

Isoproterenol (ISO) overactivation of β-adrenergic receptors in cardiomyocytes triggers pathological processes including increased myocardial oxygen consumption, oxidative stress, and fibrosis. These lead to progressive extracellular matrix expansion, impaired contractility, and ultimately heart failure. Micropeptide APPM binds to its interacting partner OGDH and activates OGDH complex (OGDHc) enzymatic activity. This enhances TCA cycle flux, boosting mitochondrial production of reducing equivalents (NADH) and ATP. Through these metabolic effects, APPM:

- Regulates mitochondrial membrane potential
- Maintains mitochondrial morphology
- Improves cellular energy supply
- Ultimately ameliorates heart failure progression

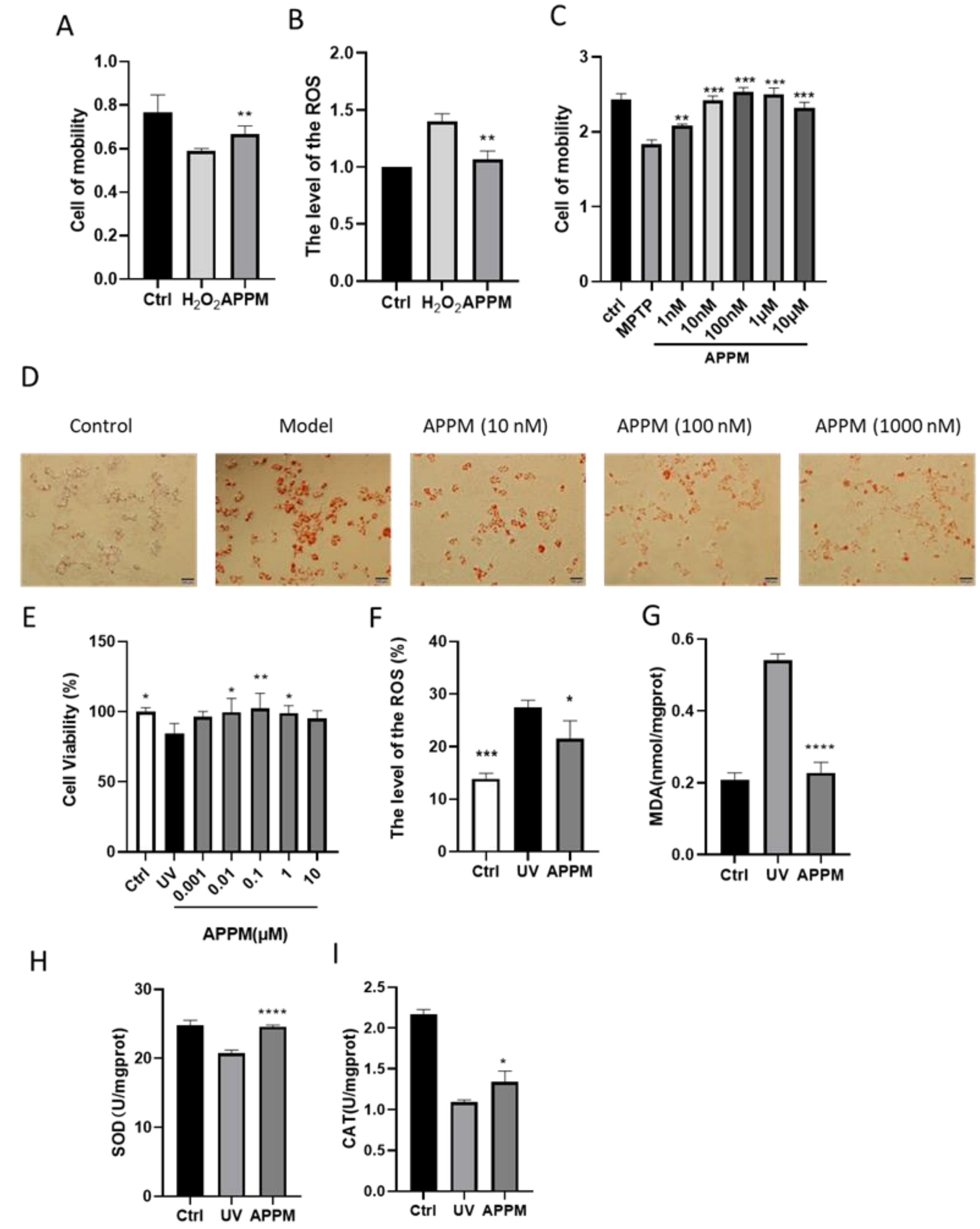


**Figure S9. The effect of APPM on disorders underlying mitochondrial bioenergetic deficiencies**

**(A)** APPM (10nM) alleviates the inhibitory effect of $H_2O_2$ on cell proliferation.

**(B)** APPM (10nM) reduces intracellular ROS levels.

**(C)** APPM (10nM) mitigates the suppressive effect of MPTP on neuronal cell proliferation.

**(D)** APPM suppresses lipid droplet formation in AML-12 cells induced by OA and PA.

**(E-I)** The effects of APPM on UV-induced apoptosis, and oxidative stress, **(E)** cell viability, **(F)** ROS, **(G)** MDA, **(H)** SOD, **(I)** CAT

Data are presented as mean ± SEM (n=3). Statistical analysis was performed using GraphPad Prism 8.0 (unpaired t-test). *P<0.05, **P<0.01 vs control group; #P<0.05 vs model group.
